# MicroRNAs are necessary for the emergence of Purkinje cell identity

**DOI:** 10.1101/2023.09.28.560023

**Authors:** Norjin Zolboot, Yao Xiao, Jessica X. Du, Marwan M. Ghanem, Su Yeun Choi, Miranda J. Junn, Federico Zampa, Zeyi Huang, Ian J. MacRae, Giordano Lippi

## Abstract

Brain computations are dictated by the unique morphology and connectivity of neuronal subtypes, features established by closely timed developmental events. MicroRNAs (miRNAs) are critical for brain development, but current technologies lack the spatiotemporal resolution to determine how miRNAs instruct the steps leading to subtype identity. Here, we developed new tools to tackle this major gap. Fast and reversible miRNA loss-of-function revealed that miRNAs are necessary for cerebellar Purkinje cell (PC) differentiation, which previously appeared miRNA-independent, and resolved distinct miRNA critical windows in PC dendritogenesis and climbing fiber synaptogenesis, key determinants of PC identity. To identify underlying mechanisms, we generated a mouse model, which enables precise mapping of miRNAs and their targets in rare cell types. With PC-specific maps, we found that the PC-enriched miR-206 drives exuberant dendritogenesis and modulates synaptogenesis. Our results showcase vastly improved approaches for dissecting miRNA function and reveal that many critical miRNA mechanisms remain largely unexplored.

**Highlights:**

- Fast miRNA loss-of-function with T6B impairs postnatal Purkinje cell development
- Reversible T6B reveals critical miRNA windows for dendritogenesis and synaptogenesis
- Conditional Spy3-Ago2 mouse line enables miRNA-target network mapping in rare cells
- Purkinje cell-enriched miR-206 regulates its unique dendritic and synaptic morphology

## Introduction

MicroRNAs (miRNAs) are short noncoding RNAs that mediate post-transcriptional repression of their mRNA targets. Accumulating evidence supports the idea that higher organismal complexity is increasingly dependent on miRNA function. While knocking out individual miRNAs in nematodes rarely leads to measurable phenotypes^1,2^, it has progressively more severe consequences from flies^3^ to zebrafish^4,5^ to mice^6^. This is consistent with evolutionary data that show a rapid expansion of the miRNA repertoire in organisms composed of many cell types^7–12^. Most of these new miRNA families are enriched in the nervous system^12^, suggesting that the brain, arguably the most complex organ, might have functioned as an “evolutionary hotspot” for post-transcriptional mechanisms. Indeed, the brain expresses the broadest variety of miRNAs among all tissues^13^, many of which are developmentally regulated^14–16^ and strongly enriched in different neuronal subtypes^16–18^. Neuronal transcripts are known to have longer 3’ untranslated regions (UTRs, a major site for miRNA binding)^19–21^, which harbor conserved sites for many brain-enriched miRNAs^20,22^ that can act cooperatively to significantly enhance the magnitude of repression^23^. The emergence of these additional post-transcriptional mechanisms may have provided efficient means by which existing gene regulatory networks can be rewired during evolution to enable neuronal diversification and provide the building blocks for new circuit motifs.

Taken together, this large body of work indicates that the brain is a prominent site for extensive miRNA function. Indeed, global miRNA loss-of-function has been shown to severely impair neuronal differentiation and early brain development^24–28^. However, further examining cell type-or developmental stage-specific miRNA function has been hindered by technologies that lack the required temporal and spatial resolution to investigate closely timed events in a cell type-specific manner. Consequently, determining if and how miRNAs instruct the developmental processes that establish neuronal subtype identity, the foundation of complex brains, remains an insurmountable challenge^29^. To solve this issue, we engineered tools for rapid and reversible modulation of miRNA activity and for precise mapping of cell type-specific miRNA-target interactions (MTIs). As proof-of-principle, we applied this new toolbox to dissect the cell type-specific miRNA mechanisms instructing the emergence of cerebellar Purkinje cell (PC) identity, one of the largest and most specialized neurons in the brain.

The classic approach for examining global miRNA function in mice has been conditional knockout (cKO) of Dicer^24–28,30–36^. Dicer cKO effectively halts canonical miRNA biogenesis in targeted cells, but Dicer cKO phenotypes can require weeks or even months to present^30–32^ (Figure S1A and S1B). The delay between Dicer cKO and phenotypic onset is likely related to the observation that miRNA half-lives vary substantially, with many miRNAs remaining at functional levels weeks after Dicer cKO^30,31,37,38^. Indeed, the effects of Dicer cKO on neuronal dendritic development, which spans late embryogenesis and the first 3-4 weeks of postnatal life in mice, are observed only if Dicer is knocked out before ∼E15.5^33,34^ (Figure S1A). Thus, while Dicer cKO clearly reveals the gross phenotypic effects that accumulate over time as a result of miRNA loss, it is difficult to observe the roles of miRNAs in processes that occur in a rapid succession, such as the discrete developmental events that lead to neuronal subtype identity.

An outstanding example of the limitations of Dicer KO is its use in PC development. PCs have a protracted developmental timeline (∼E10 to ∼P28 in mice) that is essential to fully establish their unique morphology, featuring a two-dimensional, intricately elaborate dendritic arbor that defines its identity, and to form a rich network of synaptic connections^39–41^. PC-specific Dicer KO using L7^Cre^, which turns on during early postnatal life (∼P4), leads to gradual cerebellar degeneration and ataxia starting around 13 weeks of age, but has no impact on PC morphogenesis^30^. Notably, some miRNAs in Dicer cKO PCs remain at near-physiological levels after PC morphogenesis is complete^30^, raising the possibility that miRNA function in PC development are masked by slow miRNA decay. Thus, determining whether miRNAs drive neuronal subtype identity requires tools that can modulate global miRNA activity faster than the timescale at which neuronal development occurs. Here, we took advantage of the short Tnrc6b-derived peptide - T6B - that induces rapid miRNA loss-of-function without affecting miRNA biogenesis in non-neuronal cells^42–44^. Adapting T6B to neurons, we discovered that miRNAs are in fact critical for PC development. We further developed an inducible and reversible version of T6B that can be switched on and off within several hours, allowing us to clearly separate the contributions of miRNAs to PC dendritogenesis, synapse formation, and cell survival.

To illuminate the molecular mechanisms driving these developmental processes, we created a mouse line amenable to mapping MTIs in rare cell types. Biochemical approaches to purify Ago2-miRNA-target complexes have successfully mapped MTIs in abundant populations of neurons^45,46^. However, due to their technical complexity and high background, these approaches are not suitable for mapping MTIs in rare cell populations such as PCs and the many other neuronal subtypes that compose the mammalian brain. We therefore generated a mouse line with a conditional Spy3-Tag, which is small and offers near-infinite affinity for pull-downs^47^, in the endogenous Ago2 gene. Using Spy3-Ago2 pull-down and sequencing (SAPseq), we generated a comprehensive MTI map of developing PCs and identified cell type-specific miRNA-target networks that suggest post-transcriptional mechanisms leading to neuronal diversification. Indeed, we identified a PC-enriched miRNA, miR-206 that is necessary and sufficient for PC exuberant dendritogenesis, a hallmark of PC identity. These results demonstrate how the unprecedented spatiotemporal resolution of our novel toolbox can reveal how miRNAs instruct the emergence of neuronal subtype identity, an outstanding question in the field.

## Results

### The miRNA machinery is intact and functional in developing PCs

Multiple studies have shown that miRNAs are important for PC maintenance and survival^30,48^, but there is currently no evidence that miRNAs are necessary for PC development. Because inhibition of individual miRNAs disrupts dendritogenesis and synaptogenesis in excitatory pyramidal neurons (PNs)^15,49–51^, it was surprising that global miRNA function would be dispensable for developing PCs. To examine the hypothesis that miRNAs are dispensable for PC development, we first determined if the components of the miRNA-induced silencing complex (miRISC), the effector of miRNA function, are present and functional in developing PCs. The three core miRISC proteins Ago1, Ago2, and Tnrc6a are readily detected in developing cerebella by immunofluorescence with a relative enrichment in the PC soma versus the neighboring granule cell layer, which becomes more pronounced with time (Figure S1C). To determine whether miRISC is functional in PCs, we used a fluorescent miRNA sensor to measure miRNA repression activity^15,52^. We designed an mCherry-based sensor for miR-124 (Figure S1D) because miR-124 is robustly expressed in all neurons, including PCs^17,22,30,53^. We transduced wild-type PCs at P0 with either a miR-124 sensor or a control sensor in which the miRNA response element (MRE) is disrupted by mutations (Figure S1D). At P14 and P21, we observed a reduction in mCherry fluorescence with the miR-124 sensor compared to the control, indicating that miRNA function is intact in PCs (Figure S1E). We observed a similar result in L7^Cre^; Dicer^fl/fl^ mice (Figure S1E), a mouse line that shows no defects in PC postnatal development but PC degeneration at 13 weeks of age^30^, suggesting that miRISC is still functional weeks after Dicer cKO. Together, the data show that Dicer cKO is too slow to study the effects of miRNA loss-of-function with a temporal resolution matching neuronal development.

### Fast miRNA loss-of-function with T6B induces profound PC morphogenesis defects that affect behavior

To induce rapid miRNA loss-of-function in neurons, we sought to use the peptide T6B that binds Ago with high affinity^42^. We reasoned that T6B overexpression in neurons could compete with endogenous Tnrc6 and inhibit miRNA-mediated repression (Figure 1A), as previously shown in non-neuronal cells^42–44^. Specifically, a mouse strain harboring a doxycycline-inducible T6B-YFP fusion protein transgene has previously been used to examine the consequences of acutely blocking miRNA function in non-nervous tissues such as the intestines, skeletal muscles, and the heart^43^. Unfortunately, doxycycline administration induced minimal to no T6B expression in the central nervous system^43^, preventing the use of this mouse strain to assess miRNA function in the brain. To circumvent this, we engineered neuronal-specific AAVs expressing the T6B-YFP fusion protein (T6B from here on). A peptide in which the residues essential for Ago binding are mutated was used as negative control (Ctrl, Figure 1B). We first assessed T6B function in cultured cortical PNs using the miR-124 sensor and found that T6B effectively de-repressed mCherry (Figure S1F). T6B also induced a significant reduction of dendritic complexity in PNs (Figure S1G), a phenotype previously linked to miRNA loss-of-function^33,34,49^.

**Figure 1.**
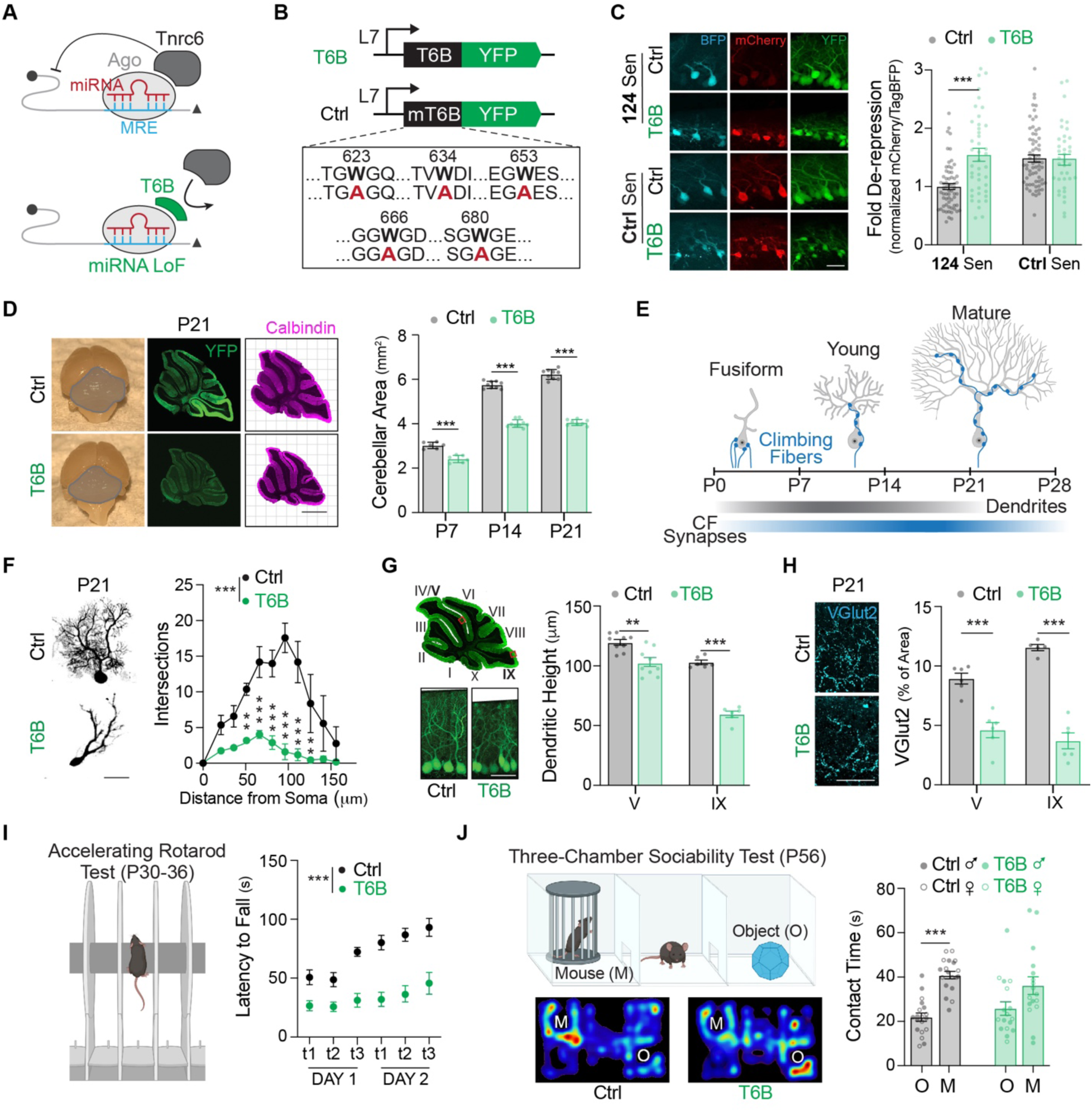
**Fast miRNA loss-of-function in PCs with T6B induces profound morphogenesis defects that affect behavior.** (A) Schematic of miRNA loss-of-function via T6B. Top: The Ago-miRNA complex recognizes the target based on sequence complementarity to the MRE (blue). Tnrc6 is recruited to mediate post-transcriptional repression. Bottom: T6B competes with Tnrc6 binding to induce miRNA loss-of-function. (B) Amino acid sequence difference between T6B and the mutated T6B (Ctrl). The five mutated tryptophan residues in red are essential for binding Ago. (C) T6B functional validation in PCs using the miR-124 sensor. T6B de-repressed mCherry to levels comparable to the Ctrl sensor in which the miR-124 MRE is fully mutated (see Figure S1D). Scale bar, 50 μm. N=37-63 cells from 3 mice. (D) Whole mount images and Calbindin-stained sagittal sections of Ctrl-and T6B-transduced cerebella collected at P21. Scale bar, 1 mm. Quantification of total cerebellar area in sagittal sections at P7, P14 and P21 show T6B-induced reduction. N=9 sections from 3 mice. (E) Schematic of the timeline of PC dendritogenesis and CF synaptogenesis. (F) Sholl analysis of PC dendritic complexity at P21. Sparse labeling was achieved by co-transducing an mRuby3-expressing AAV. T6B sharply decreased dendritic complexity. Scale bar, 50 μm. N=8 cells from 3 mice. (G) T6B decreased PC dendritic height in cerebellar lobules V and IX (see top left image for lobules location). Scale bar, 50 μm. N=9 sections from 3 mice. (H) T6B decreased CF synapses, visualized by VGlut2 staining, at P21. Scale bar, 50 μm. N=6-9 sections from 3 mice. (I) T6B-transduced mice fell sooner from the accelerating rotarod. N=18 mice. (J) The three-chamber sociability test is depicted (top left). T6B-transduced mice showed less preference for a novel mouse (M) vs a novel object (O). Bottom left: representative heat maps of mouse location. Male and female mice are depicted with full circles or border only, respectively. N=18 mice. Data are mean ± SEM. Statistics for (C), (D), (G), (H): Welch’s t-test; (F): Mixed-effects model with Šídák’s multiple comparisons test; (J): Mann-Whitney test; (I): Two-way repeated measures ANOVA; (J): Two-way ANOVA. *p ≤ 0.05; **p ≤ 0.01; ***p ≤ 0.001. See also Figure S1 and S2.

To examine miRNA roles exclusively in developing PCs, we cloned the PC-specific minimal L7 promoter^54^ upstream of T6B (Figure S2A). Co-transduction of L7-T6B and the miR- 124 sensor confirmed miRNA loss-of-function in PCs *in vivo* (Figure 1C). Contrary to L7^Cre^; Dicer^fl/fl^ mice, T6B-transduced mice had smaller cerebella than controls, which was noticeable from P7 (Figures 1D and S2B). T6B-transduced PCs had shorter and less complex dendritic arbors and smaller somatic areas (Figure 1F, 1G and S2C), a phenotype that resembles miRNA loss-of-function in PNs^34^, indicating that regulating dendritic and somatic growth is likely a common function for miRNAs in neurons. Further, T6B induced smaller granule cell layer (GCL) area (Figure S2D) and reduced synaptic input from climbing fibers (CF) of the inferior olivary nucleus onto PCs (Figure 1E and 1H), which is critical for motor coordination and learning^55^.

Compared to L7^Cre^; Dicer^fl/fl^ mice, where Dicer cKO led to PC death at P90^30^, T6B induced PC loss much earlier. At P28, we observed significant PC loss in lobules V and IX (Figure S2E). Increased cleaved caspase-3 in T6B-transduced PCs suggests that cell loss is induced by apoptosis (Figure S2F). Intracerebroventricular injection of AAV appears to target cerebellar lobule IX more efficiently, leading to stronger phenotypes compared to other lobules (Figure S2G). T6B-induced cell loss becomes progressively more severe and by P90, most PCs in lobule V are also lost (Figures S2H). Consistent with the severity of the cellular phenotype, T6B-transduced mice showed defects in behaviors classically linked to PC function, such as impaired motor coordination, altered sociability, hyperactivity in the open field, and reduced stride length (Figure 1I, 1J and S2I-S2L). Similar behavioral deficits have been observed with genetic perturbations that alter PC morphology and their synaptic input^56–58^. These phenotypes are core features of ataxia and autism spectrum disorder, conditions strongly linked to impairments in PC development and function^41^. Intriguingly, miRNAs have numerous roles in the pathogenesis of spinocerebellar ataxia^59^, which uniquely affects PCs compared to other neurons. Together, the cellular and behavioral phenotypes observed in T6B-transduced mice indicate that miRNA function is critical to normal PC development, contrary to what was shown in previous Dicer cKO studies. Our findings with T6B-induced rapid miRNA loss-of-function suggest that results of Dicer cKO experiments have significantly underestimated the importance of miRNAs during neuronal development.

### A reversible T6B peptide decouples cell death from morphogenesis defects

The cell death induced by T6B makes it difficult to discern whether the observed changes are a developmental phenotype or simply a by-product of imminent PC apoptosis. To distinguish between these possibilities, we envisioned an inducible and reversible T6B system to determine if transient miRNA loss-of-function would differentially affect development versus cell survival. The doxycycline-inducible tet-on system (used in the T6B transgenic mouse strain) provides a way to transiently block miRNA function, where doxycycline induces the expression of a transactivator, which in turn activates T6B transcription^43^. However, this strategy is relatively slow, requiring several days to reach full T6B expression levels after doxycycline administration and several more days to clear T6B after doxycycline removal^43^. Instead, directly targeting T6B would provide rapid and tunable regulation of protein levels. To modulate miRNA activity on a time scale better matched for neuronal development, we engineered a rapidly inducible and reversible T6B construct. T6B was fused to the dihydrofolate reductase destabilizing domain (DD) from *E. coli*, which directs fast proteasomal degradation of newly synthesized peptides^60^. The degradation of DD-T6B can be transiently blocked by Trimethoprim (TMP), a small molecule with high affinity for DD^60–62^ (Figure 2A). In cultured PNs, TMP-stabilized DD-T6B de-repressed the miR-124 sensor and reduced dendritic complexity to degrees comparable to T6B but had no measurable effect in the absence of TMP (Figures 2B and 2C).

**Figure 2.**
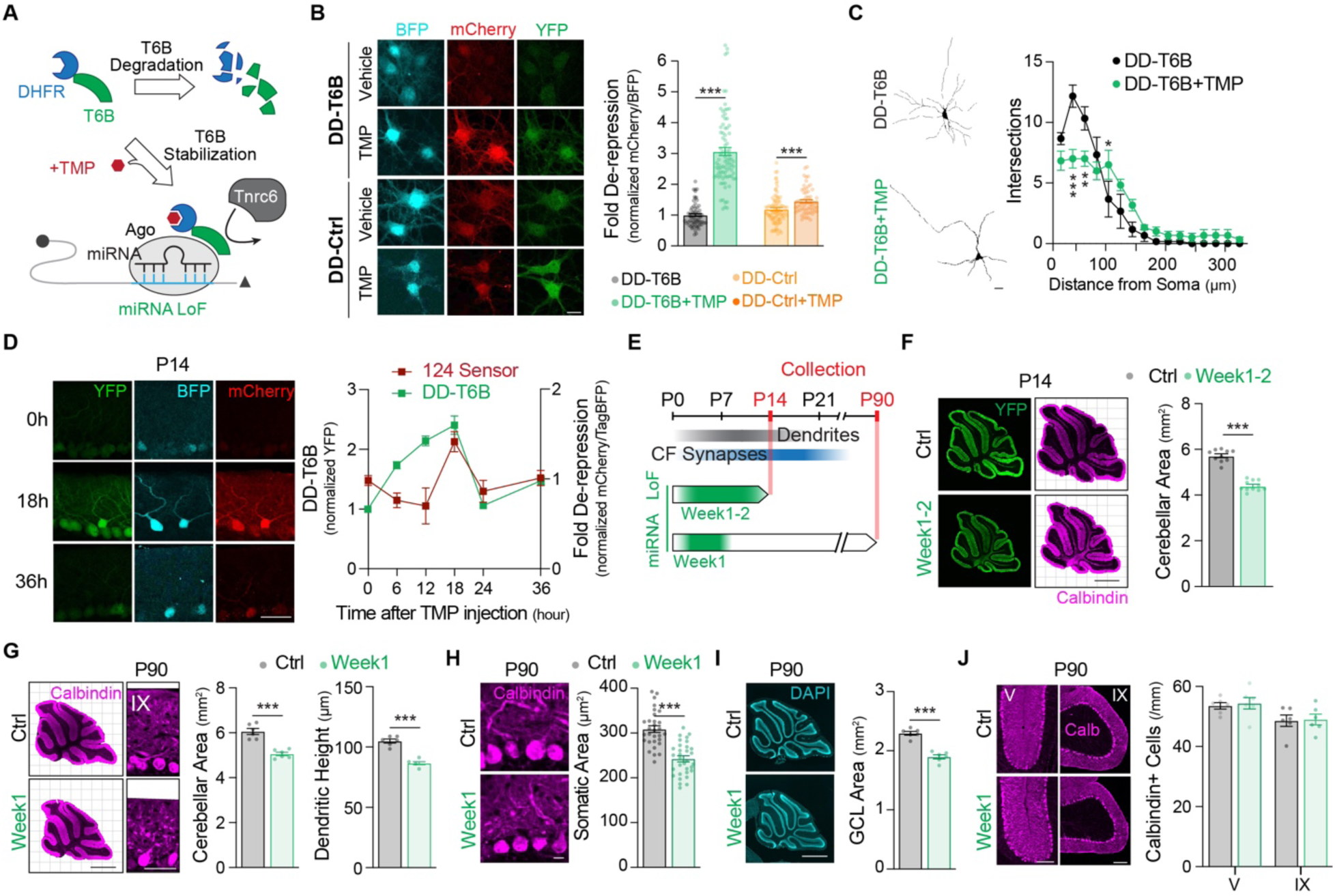
**A reversible T6B (DD-T6B) decouples cell death from morphogenesis defects** (A) Schematic of DD-T6B function. The DHFR destabilizing domain (DD) fused to T6B induces degradation. TMP stabilizes DHFR allowing DD-T6B to rapidly accumulate and induce miRNA loss-of-function. (B) Functional validation of DD-T6B in cultured cortical PNs using the miR-124 sensor. TMP-induced DD-T6B de-repressed mCherry at DIV11. We observed de-repression of the miR-124 sensor in the presence of TMP-induced DD-Ctrl. This suggests a mild effect of TMP on mCherry levels. However, it is important to notice that the scale of the de-repression of mCherry with DD-Ctrl after TMP is small (20%) when compared to DD-T6B after TMP (300%). Scale bar, 10 μm. N=61-86 cells. (C) Sholl analysis of cultured cortical PNs at DIV11. TMP-induced DD-T6B decreased dendritic complexity. Scale bar, 10 μm. N=8 cells. (D) DD-T6B induction time course and functional validation using the miR-124 sensor in PCs at P14. DD-T6B peaks at 18 hours after TMP injection and induces mCherry de-repression. Scale bar, 50 μm. N=14-20 cells. (E) Outline of the DD-T6B experiments in panels F-J. miRNA loss-of-function (LoF) via DD-T6B was induced daily for the first two postnatal weeks (Week1-2, green segment) or for the first week (Week1, green segment). Mice were collected at P14 or P90 (red lines). (F) Calbindin-stained sagittal slices at P14 show that DD-T6B induction during the first two weeks reduced cerebellar area compared to Ctrl (from mice that were administered PBS instead of TMP). Scale bar, 1 mm. N=9 sections from 3 mice. (G) Calbindin-stained sagittal slices at P90 show that DD-T6B induction during week 1 caused an irreversible defect in cerebellar area and lobule IX PC dendritic height. Scale bar, 1mm for area; 50 μm for dendritic height. N=6 sections from 3 mice. (H) DD-T6B induction during week 1 irreversibly reduced PC soma area. Scale bar, 10 μm. N=30 cells from 3 mice. (I) DAPI-stained sagittal slices at P90 show that DD-T6B induction during week 1 irreversibly reduced GCL area. Scale bar, 1mm. N=6 sections from 3 mice. (J) DD-T6B induction during week 1 did not cause PC loss at P90, effectively decoupling developmental defects from apoptosis. Scale bar, 100 μm. N=6 sections from 3 mice. Data are mean ± SEM. Statistics for (A), (B), (F), (G), (H), (I) and (J): Welch’s t-test; (C): Mixed-effects model with Šídák’s multiple comparisons test. **p ≤ 0.01; ***p ≤ 0.001. See also Figure S3.

To assess the performance of DD-T6B in PCs, we cloned DD-T6B downstream of the L7 promoter. DD-T6B had no effect on cerebellar area or PC dendritic height in the absence of TMP (Figure S3A). After a single TMP injection, DD-T6B reached peak expression within 24 hours, and was undetectable 36 to 96 hours later depending on the developmental stage (Figures 2D and S3B). *In vivo* miR-124 sensor data show that miRNA activity is turned off and back on in a much narrower timeframe (Figure 2D). The miR-124 sensor is acutely de-repressed once DD-T6B reaches peak expression (18 hours after TMP injection) and immediately re-repressed once DD-T6B levels decrease, suggesting that this strategy enables blocking miRNA function within a very tight temporal window (Figure 2D). To determine whether stabilized DD-T6B can recapitulate T6B phenotypes, we induced DD-T6B for 2 weeks with daily TMP injections (Figure 2E). We found that TMP-induced DD-T6B reduced cerebellar area (Figures 2F) to a similar extent as T6B (Figure 1D and S2B).

We next asked if miRNAs are necessary for PC development independently of cell survival by transiently inducing miRNA loss-of-function during the first postnatal week. TMP was injected daily starting at P1 and stopping at P7 (Week 1). Based on the DD-T6B induction time course (Figure S3B), we expect miRNA function to be lost during the first week and fully restored by P9 (Figure 2E). Examining the cerebella at P90 revealed that miRNA loss-of-function during week 1 decreased cerebellar area, dendritic height, somatic area, and GCL area (Figure 2G-2I). However, we did not observe any cell loss in lobules V or IX (Figures 2J), indicating that miRNA function is in fact critical to normal PC development and the phenotype is not a by-product of apoptosis. Thus, DD-T6B effectively decouples cell death, which inevitably occurs if miRNA loss-of-function persists^25,27,28,30,36^, from morphogenesis defects.

### DD-T6B reveals critical miRNA windows for PC dendritogenesis and CF synaptogenesis

PC morphogenesis spans ∼4 postnatal weeks^39,40^, in which overlapping formative events instruct dendritic development and circuit integration. PC morphology progresses from fusiform to having highly arborized dendrites, reaching maximal complexity at ∼P20^39^. Innervation by CFs occurs during a similar time frame. Initially, each PC soma is innervated by multiple CFs. Then a single CF is selectively strengthened and by P14, only one CF remains, which then climbs the dendritic arbor^63^ (Figure 1E).

To dissect the roles of miRNAs in PC dendritogenesis and CF synaptogenesis, we induced transient miRNA loss-of-function using DD-T6B at week 1, 2, or 3 and analyzed cerebella at P28, so that all conditions have at least one week of restored miRNA function before analysis (Figure 3A). Week 4 was used as a control, because PC morphogenesis is mostly completed by end of week 3^39^. miRNA loss-of-function during week 1 and 2, but not 3 or 4, reduced cerebellar area and PC dendritic height at P28 (Figures 3B and 3C), but to a lesser extent than constitutively expressed T6B (Figures 1D, 1G). Somatic area and GCL area were also reduced when miRNA function was disrupted during week 1 and 2, indicating that miRNA function is critical for these processes during this timeframe (Figure 3D and 3E). Interestingly, only miRNA loss-of-function during week 3 induced a consistent decrease in the density of synaptic puncta from CFs on primary, secondary, and tertiary PC dendrites (Figures 3F and S3C). CF translocation to its maximal height was not affected in any of the conditions (Figure S3C). These results reveal separate miRNA-mediated mechanisms that instruct PC dendritogenesis and CF synaptogenesis. Notably, a stunted dendritic arbor (achieved by miRNA loss-of-function in week 1 and 2) does not affect CF translocation, indicating that, during week 3 miRNAs regulate signaling in the post-synaptic cell to instruct proper CF innervation. We conclude that week 1 and week 3 are critical miRNA developmental windows for dendritogenesis and CF synaptogenesis, respectively. DD-T6B offers an approach for assessing miRNA function with greater temporal precision and has allowed us to decouple two developmental events that are overlapping in time.

**Figure 3.**
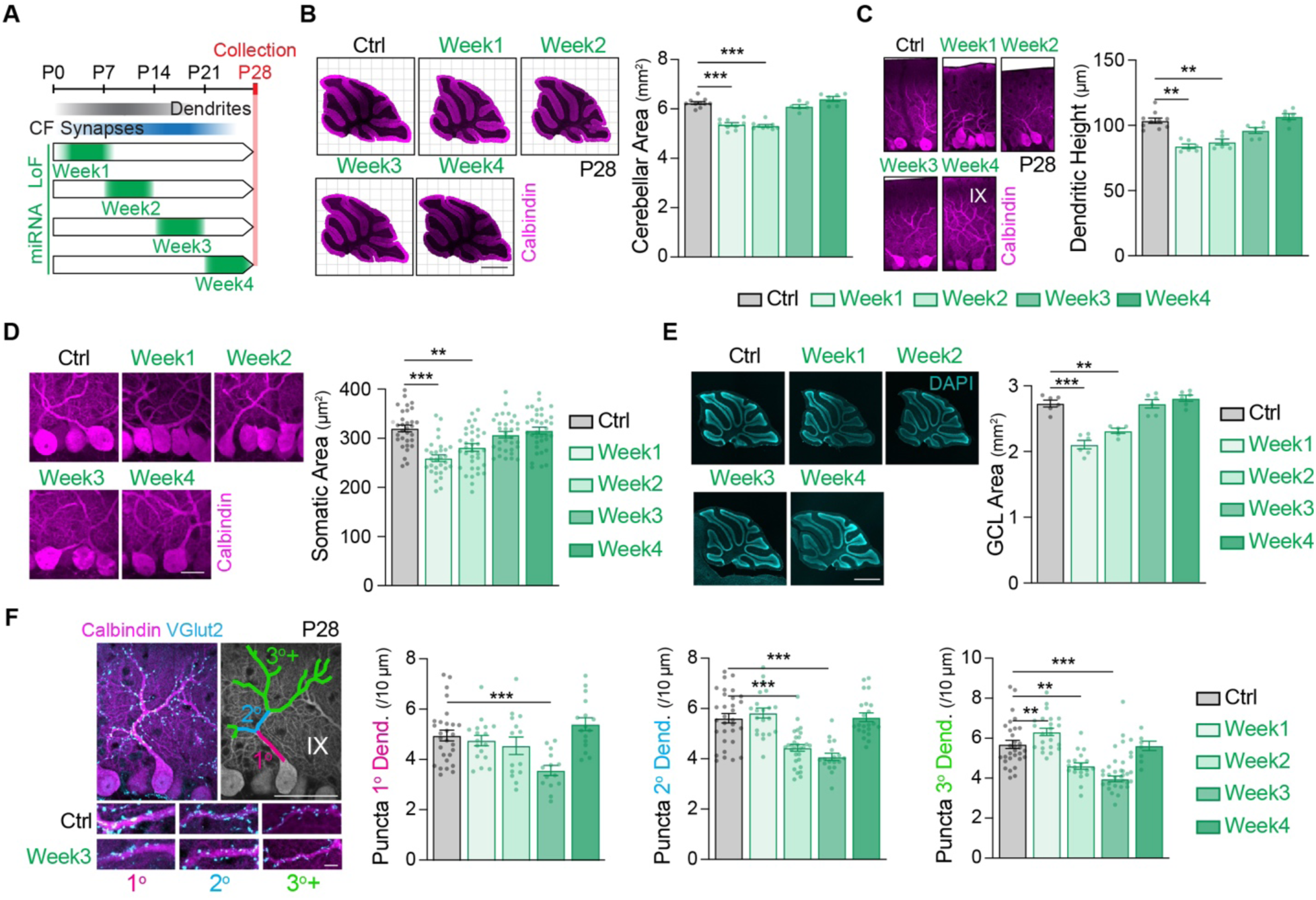
**DD-T6B reveals critical miRNA windows for PC dendritogenesis and CF synaptogenesis** (A) Outline of the DD-T6B experiments in this figure. miRNA loss-of-function (LoF) via DD-T6B was induced for one week (green segment) in either week 1, week 2, week 3, or week 4. In each case (except week 4) miRNA function was restored for at least one week to allow for recovery of a possible developmental delay. Mice were collected at P28 (red line). (B and C) Calbindin-stained sagittal slices at P28 show that DD-T6B induction during week 1 or 2 irreversibly reduced cerebellar area and PC dendritic height. Scale bar, 1 mm for area; 50 μm for dendritic height. N=6 sections from 3 mice. (D) DD-T6B induction during week 1 and week 2 reduced PC soma area. Scale bar, 10 μm. N=30 cells from 3 mice. (E) DAPI-stained sagittal slices at P28 show that DD-T6B induction during week 1 and week 2 reduced GCL area. Scale bar, 1 mm. N=6 sections from 3 mice. (F) Immunostaining and quantification of VGlut2 puncta density in PCs at P28. Primary (fuchsia), secondary (blue) and tertiary (green) dendrites were quantified separately. DD-T6B induction during week 3 consistently reduced VGlut2 puncta on all dendrites. Induction during week 1 caused an increase in puncta on tertiary dendrites. Scale bar, 50 μm. N=14-32 dendrites from 3 mice. Data are mean ± SEM. Statistics for (B), (C), (D), (E) and (F): Two-way ANOVA. **p ≤ 0.01; ***p ≤ 0.001. See also Figure S3.

### SAPseq identifies miRNA-target interactions with minimal background and perturbation of miRNA function

Our data show that miRNAs are necessary to drive exuberant dendritogenesis and CF synaptogenesis. To determine the mechanisms driving these unique features, it is essential to first map the post-transcriptional gene regulatory network underlying these developmental processes. miRNA-target networks are highly complex and notoriously difficult to map. This challenge is exacerbated in rare cells, such as the many neuronal subtypes that compose the mammalian brain. Mapping of neuronal subtype-specific MTIs has previously been achieved from abundant neuronal populations using either crosslinking-immunoprecipitation (CLIPseq) of a Flag-tagged conditional Ago2 transgene^45,46^ or Halo-enhanced Ago2 pull-down (HEAPseq) from a Halo-Ago2 conditional knock-in allele^64^. However, none of these techniques has been used in rare cell types, such as PCs that make up less than 1% of cells in the cerebellum^65^. To map MTIs in PCs, we first tried Ago2 CLIPseq using a commercially available Myc-Ago2 transgenic mouse line (tAgo2)^17^. However, this yielded poor signal-to-noise with a peak distribution inconsistent with the known bias for mRNA 3’ UTR^16,22^ (Figure S4A), indicating that the antibody-based IP approach is poorly suited for rare neuronal populations such as PCs. We next sought to try HEAPseq, which achieves superior signal-to-noise by taking advantage of a covalent bond that HaloTag forms with its immobilized ligand^64^. Additionally, the Halo-Ago2 is tagged at the endogenous Ago2 locus and thus does not require overexpression from a transgene. Considering the complex patterns of Ago2 parental imprinting in the brain^66,67^, and that PCs make up less than 1% of cells in the cerebellum^64^, we sought to perform the pull-down from homozygous mice. However, homozygous Halo-Ago2 mice (Ago2^Halo/Halo^) were not viable (Figure S4B), suggesting that the insertion of the large (>1 kb) HaloTag conditional cassette in the Ago2 gene promoter region, or appending the HaloTag domain to the Ago2 protein, is deleterious to some aspects of Ago2 function. Moreover, considering that even a single amino acid mutation in Ago2 can lead to global neurodevelopmental delay^68^, the potential disruption a bulky HaloTag (33kDa) could have on neuronal Ago2 function was a cause for concern.

To circumvent these issues, we created a tagged Ago2 mouse and a mouse embryonic stem cell (mESC) line designed for minimal perturbation of miRISC function. We used a cleavable SpyTag3^47^, which rapidly forms a covalent bond with its ligand SpyCatcher3 and is ten times smaller than HaloTag (3 vs 33 kDa). To minimize any perturbation on Ago2 expression and protein structure, constitutive Spy3-Ago2 mice and mESC lines were created by inserting the SpyTag3 coding sequence into exon 2 of the Ago2 gene, which is separated from the promoter region by a 38 kb intron and encodes a small unstructured region of Ago2 that is the only part of the protein sequence that is not perfectly conserved between mice and humans (Figures S4C and S4D). Ago2^Spy3/Spy3^ mice were born at Mendelian frequencies (Figure S4B). Ago2 levels were reduced in Ago2^Spy3/+^ and Ago2^Spy3/Spy3^ mouse brains (Figure S5A), but repression of the miR- 124 sensor was comparable to wild-type mice (Figure S5B), indicating that Ago2 function is unaffected, at least for abundant miRNAs. To test whether Spy3-Ago2 can be used to map MTIs, we performed SAPseq (Figure 4A), a method developed based on HEAPseq^64^, using mESCs (Figure S5C). The miRNA and mRNA target libraries that we generated with SAPseq were highly reproducible between biological replicates (Figure S5D-S5F). From the target library we were able to call 3292 reproducible peaks with the CLIPanalyzer package^63^, more than 40% of which aligned to the 3’ UTR (Figure 4B). We observed a positive correlation between the relative abundance of the miRNAs that were pulled down and the number of peaks containing seed-matched sequences for those miRNAs, indicating that SAPseq captures robust MTIs (Figure 4C). Moreover, the SAPseq MTI map correlated well with a mESCs HEAPseq dataset^64^ (Figures 4D-4F), demonstrating that it has similar performance to benchmark techniques, while being considerably less disruptive to Ago2 function.

**Figure 4.**
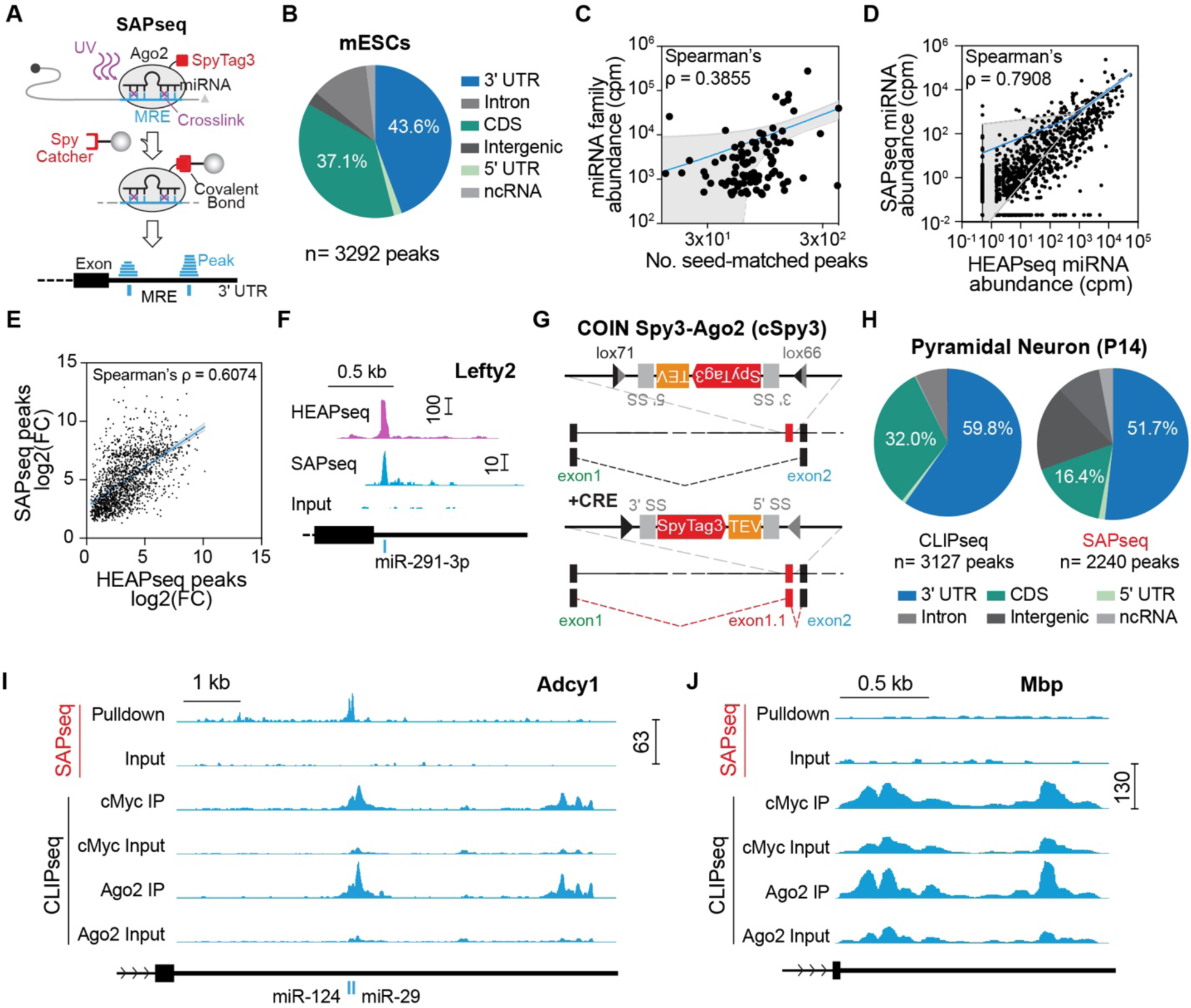
**SAPseq identifies miRNA-target interactions with minimal background and perturbation of miRNA function** (A) Schematic of SAPseq. Top: Ago2 is tagged with SpyTag3 (red). The Ago2-miRNA-target complex is crosslinked with UV. Middle: RNAse digests the target fragments not protected by Ago2 (dotted gray line). SpyCatcher3 beads are used to pull down Ago2. Bottom: Pulled-down target fragments and miRNAs are sequenced. The target fragments (blue) form peaks where MTIs are frequent, often in the 3’ UTR. (B) Frequency across genomic annotations of the peaks mapped in the E14 mESCs. More than 40% of the peaks mapped to the 3’ UTR. (C) The number of SAPseq peaks with 7-mer or 8-mer seed matches to the top 100 expressed miRNA families versus the abundance of the corresponding miRNA family. (D) Pairwise comparison of SAPseq and HEAPseq miRNA libraries generated from mESCs show positive correlation. (E) Pairwise comparison of SAPseq and HEAPseq target libraries generated from mESCs show positive correlation. Per peak log2FC over input was compared. (F) Genome browser view of peaks identified in the Lefty2 3’ UTR. Both HEAPseq and SAPseq identify a peak that is assigned to miR-291-3p. (G) Schematic of the conditional by inversion (COIN) strategy used to generate the cSpy3-Ago2 line. (H) Frequency across genomic annotations of the peaks mapped from P14 cortical PNs with SAPseq and CLIPseq. (I) Genome browser view of peaks identified in the Adcy1 3’ UTRs in P14 PNs using SAPseq from VGlut2^Cre^ ^+/-^; Ago2^cSpy3/+^; cMyc CLIPseq (PN-specific) and Ago2 CLIPseq (not cell type-specific) from VGlut2^Cre^ ^+/-^; tAgo2^+/-^ mice. SAPseq (top), cMyc and Ago2 CLIPseq (bottom) show similar peaks. SAPseq peaks are sharper and fewer. Predicted MREs are shown below. (J) Peaks identified in the Mbp 3’ UTRs. Mbp is a major constituent of myelin sheaths made by oligodendrocytes, so it should not be expressed in neurons. SAPseq shows no peaks (top) while peaks appear in cMyc and Ago2 CLIPseq (bottom), likely due to Ago2 redistribution in solution during CLIPseq. See also Figure S4 and S5.

### Conditional Spy3-Ago2 maps PC-specific miRNA-target interactions

The Spy3-Ago2 mouse line allows precise mapping of MTIs from endogenous Ago2. However, because the allele is constitutively expressed in every cell of the mouse, it cannot resolve MTIs from tissues with high complexity of cell types, such as the brain. Hence, to map cell type-specific MTIs with SAPseq, we generated a conditional by inversion^69^ (COIN) Spy3-Ago2 mouse line (cSpy3-Ago2, Figure 4G). To minimize the impact of the COIN cassette on Ago2 gene expression, the SpyTag3 coding sequence, flanked by splice and lox sites, was inserted in an inverted orientation into a poorly conserved region of the first Ago2 intron (Figure S4E). Upon Cre recombination, the SpyTag3 exon and the flanking splice sites are inverted, leading to the inclusion of SpyTag3 in the mature Ago2 mRNA (Figures 4G and S5G). Ago2^cSpy3/cSpy3^ mice were born at Mendelian frequencies (Figure S4B).

To validate the cSpy3-Ago2 mouse line, we first crossed Ago2^cSpy3/cSpy3^ to VGlut2^Cre^ mice, a Cre line that targets PNs, which constitute ∼60% of the cells in the cortex^70,71^. Endogenous Ago2 levels were not affected in VGlut2^Cre^ ^+/-^; Ago2^cSpy3/+^ mice (Figure S5H). In contrast, VGlut2^Cre^; Ago2^Halo/+^ cortices had reduced Ago2 levels (Figure S5H), suggesting that the cSpy3-Ago2 mouse line is less disruptive to miRNA function. Using SpyCatcher3, we were able to pull down Spy3-Ago2 from VGlut2^Cre+/-^; Ago2^cSpy3/+^ cortex (Figure S5I). To determine if cSpy3-Ago2 mice can be used to map cell type-specific MTIs, we compared SAPseq data from VGlut2^Cre^ ^+/-^; Ago2^cSpy3/+^cortices to Ago2 CLIPseq data from VGlut2^Cre^ ^+/-^; tAgo2^+/-^ cortices. The peak distribution obtained with both techniques was similar (Figure 4H). Moreover, SAPseq peaks were sharper (smaller footprint), consistent with improved signal-to-noise (Figure 4I). Specifically, we did not observe peaks from oligodendrocyte-enriched genes that we detected with CLIPseq, which possibly occur due to redistribution of Ago2 in solution (Figure 4J), a known confounding factor in CLIPseq^72^.

Finally, to map PC-specific MTIs, we transduced Ago2^cSpy3/cSpy3^ mice with an AAV expressing Cre under the PC-specific promoter L7 and performed SAPseq at P7 and P15. Contrary to Ago2 CLIPseq, SAPseq peaks were sharp and enriched in 3’ UTR sequences (Figures 5A and 5B). Comparing PC and PN maps, we found transcripts known for their expression in both PCs and PNs that had similar peak profiles (Figure 5C). We identified peaks in PC-specific transcripts and developmental changes in MTIs between P7 and P15 (Figures 5B and 5D), indicating that PC-specific mapping of MTIs was successful. For further analysis, we only considered peaks in the 3’ UTR, the canonical site of MTIs. GO Term analysis of the miRNA targets at both ages showed many terms in common, but P15 had stronger enrichment for axonal development and synapse assembly, indicating a developmental shift in miRNA function (Figure 5E). We assigned miRNAs to the 3’ UTR peaks and built bipartite miRNA-target networks^16^ to identify the most active miRNAs and the mRNAs most heavily targeted (Figure 5F). This analysis revealed several abundantly expressed pan-neuronal miRNAs (i.e., miR-9, miR-124, miR-138) as well as miRNAs that are understudied in neurons (i.e., miR-206). Heavily targeted transcripts included those known for their selective expression in PCs within the cerebellum (Adcy1) as well as transcripts known to regulate PC dendritic growth (Shank3^73^ and Dab2ip^74^). These results demonstrate that the unprecedented spatial resolution of SAPseq can accurately map miRNA-target networks in rare cell types, a long-standing unmet need in the field.

**Figure 5.**
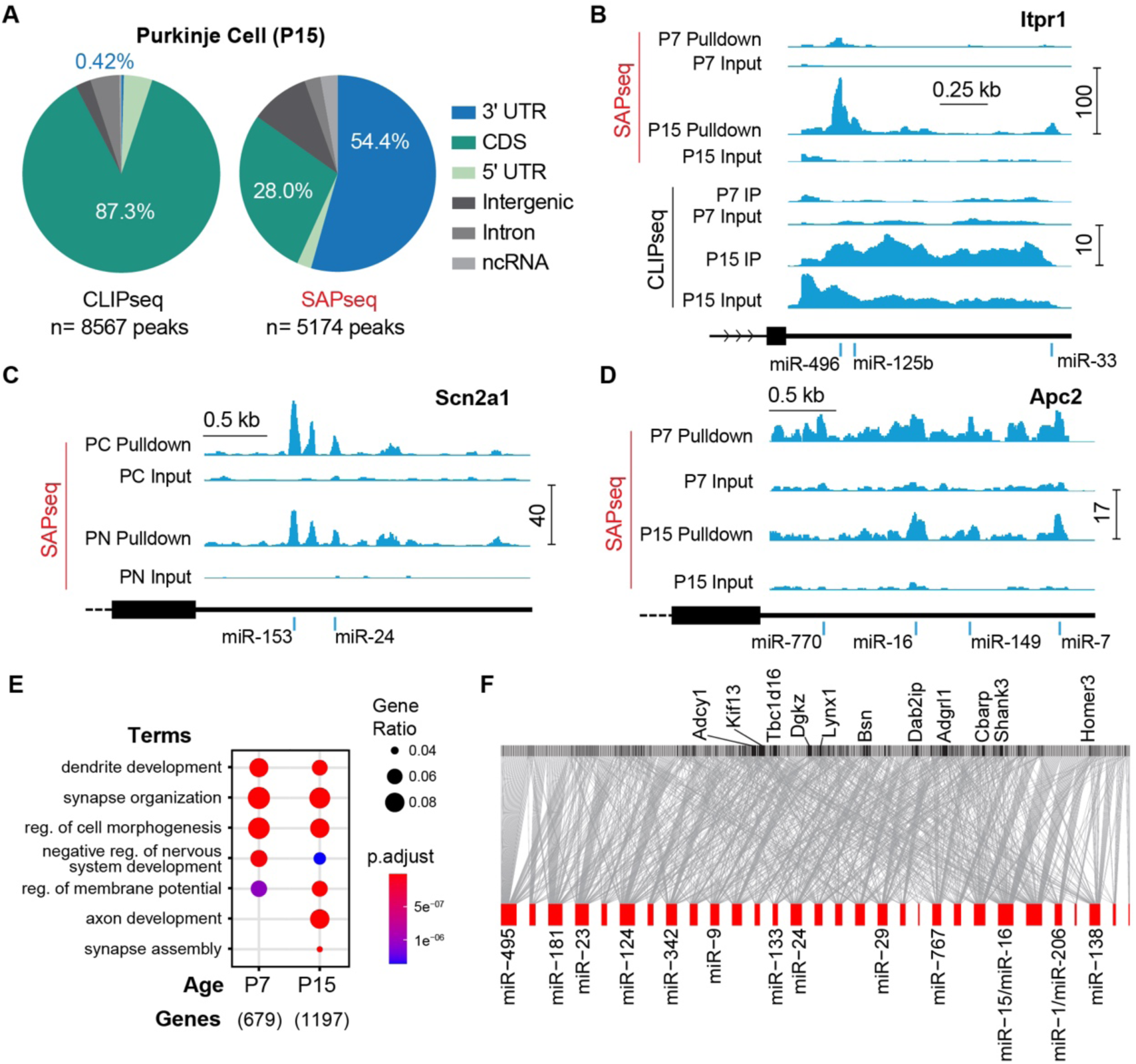
**Conditional Spy3-Ago2 maps PC-specific miRNA-target interactions** (A) Frequency across genomic annotations of the peaks mapped from P15 PC SAPseq and cMyc CLIPseq. (B) Peaks identified in the Itpr1 (PC-specific) 3’ UTRs at P7 and P15. Top: sharp peaks over low input with SAPseq. Developmental increase from P7 to P15. Bottom: small peaks with cMyc CLIPseq (notice that the scale for CLIPseq peak size is 10 vs 100 for SAPseq) over noisy input. Predicted MREs are shown below. (C) Peaks identified in the Scn2a1 3’ UTRs in P15 PCs and P14 PNs. (D) Peaks identified in the Apc2 (PC-specific) 3’ UTRs at P7 and P15 PCs using SAPseq. Developmental decrease from P7 to P15. (E) GO Term analysis of miRNA targets in PCs at P7 and P15. The terms differentially enriched at either age are due to differential miRNA targeting and not changes in transcript levels. Also, the enrichment does not reflect subcellular localization of Ago2, as Ago2 is mostly enriched in PC somas at this stage (Figure S1C). (F) Bipartite network of the top 30 most abundant miRNAs and their assigned targets in P15 PCs.

### PC-enriched miR-206 is sufficient and necessary for PC dendritogenesis

PCs have unique morphological and connectivity features, such as their highly branched dendritic arbor and innervation by CFs. Having mapped MTIs in developing PCs provided the opportunity to ask if PC-specific miRNA mechanisms drive these features. We first sought to determine a measure of how strongly each target is bound by miRNAs. For each target, the reads of each 3’ UTR peak were summed and normalized by library depth and 3’ UTR length to obtain a SAPseq RPKM (reads per kilobase of transcript per million reads mapped) value. We then plotted the SAPseq RPKM values against a P14 PC RNAseq dataset^75^ and found a positive correlation (Figure S6A). Then, we calculated and assigned a “SAP score” for each target based on distance from the line of fit^76^ (Figure S6A). A positive SAP score indicates that a target receives stronger Ago2-miRNA engagement than predicted by its transcript levels. To identify cell type-specific miRNA-target networks, we compared the SAP score of PC vs PN targets and found that many mRNAs are similarly targeted in both cell types, but a subset is differentially targeted (Figure 6A). GO Term analysis revealed that transcripts differentially targeted in PCs are enriched for the “dendrite development” term (Figure 6B, e.g., Shank3, Dab2ip), suggesting that dendritogenesis in PCs requires additional regulation by miRNAs. Interestingly, the term “synapse organization” is enriched in the common targets, an example of a post-transcriptional program that is important for both cell types. It is also enriched in the transcripts more targeted in PCs or PNs, suggesting that cell type-specific miRNA-target networks contribute to their unique connectivity.

**Figure 6.**
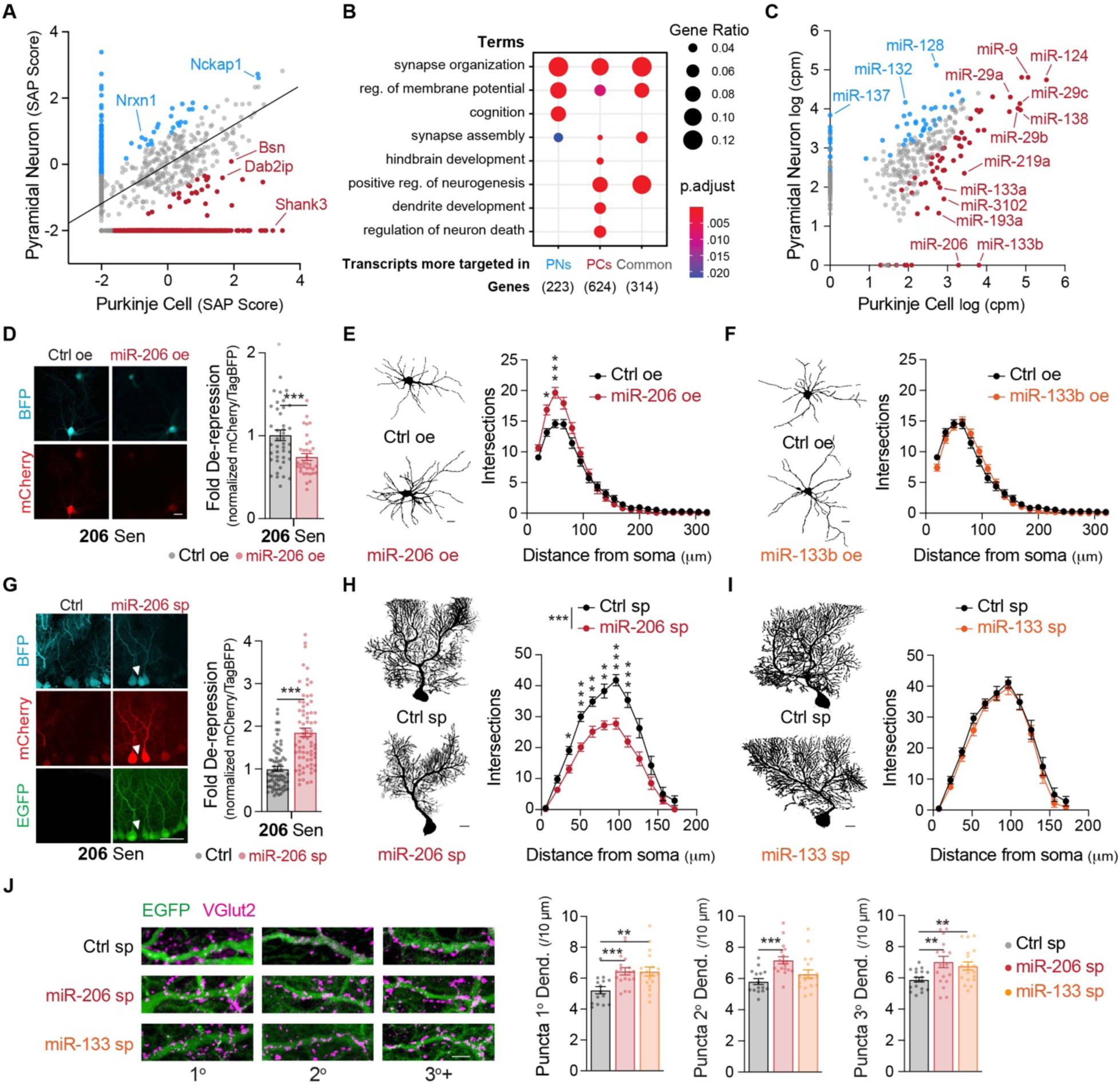
**PC-enriched miR-206 is sufficient and necessary for PC dendritogenesis.** (A) Comparison of SAP scores in PNs vs PCs. The line represents best-fit Deming regression. Transcripts that are more targeted in PCs or PNs are in red or blue, respectively. (B) GO Term analysis of the transcripts more targeted in PCs, in PNs, or similarly targeted in both (Common). (C) Normalized miRNA abundance plotted for PNs vs PCs (log scale). PC-enriched miRNAs are in red. PN-enriched miRNAs are in blue. (D) Validation of miR-206 overexpression (oe) in PNs with a miR-206 sensor (206 Sen). (E and F) Sholl analysis in DIV14 cultured cortical PNs that were co-transfected with a plasmid expressing iRFP670. miR-206 overexpression increases proximal dendritic complexity. Scale bar, 10 μm. N=31-33 cells. (G) Validation of the miR-206 sponge with the 206 Sen in P28 PCs, showing de-repression of mCherry. Note that mCherry remains repressed in PCs not expressing the sponge (white arrow). The Ctrl here are littermate wildtype mice (L7^Cre^ is a BAC line) transduced with 206 Sen. Scale bar, 50 μm. N=88-160 cells from 3 mice. (H and I) Sholl analysis of PCs at P28. miR-206 loss-of-function decreases PC dendritic complexity. Scale bar, 10 μm. N=18-23 cells from 3 mice. (J) miR-206 or miR-133 (the sponge is likely bound by both miR-133a and -133b) loss-of-function increases CF puncta in primary and tertiary dendrites at P28. Scale bar, 10 μm. N=30 dendrites from 3 mice. Data are mean ± SEM. Statistics for (A): Deming regression; (C): Wald test with DESeq2; (E) and (H): Mixed-effects model with Šídák’s multiple comparisons test; (D), (G) and (J): Welch’s t-test. **p ≤ 0.01; ***p ≤ 0.001. See also Figure S6.

To identify miRNAs potentially driving cell type-specific differences, we compared miRNAs pulled down with SAPseq from PCs or PNs. Most miRNAs are expressed at similar levels, but a subset is enriched in each cell type (Figure 6C). We then asked whether expressing PC-enriched miRNAs (miR-206 and miR-133b) in PNs could induce PC-like features. In cultured PNs, we found that overexpression of miR-206, but not of miR-133b (validation in Figures 6D and S6B), increased the complexity of their proximal dendrites (Figure 6E and 6F), suggesting that PC-like features are partially driven by PC-enriched miRNAs repressing targets expressed in both cell types. To determine whether miR-206 and miR-133 are necessary for unique features in PCs, we designed Cre-inducible miRNA sponges^49,50,77,78^, artificial miRNA targets containing 12 high-affinity MREs (Figures S6C-S6E). As validation, we showed that in cultured PNs, a miR-124 sponge de-repressed the miR-124 sensor (Figure S6F), and in L7^Cre^ PCs, the miR-206 and miR-133 sponges could de-repress their corresponding sensors (Figures 6G, S6G and S6H), indicating that the sponges are efficiently sequestering their associated miRNAs. In PCs, we found that sponging miR-206, but not miR-133, reduced the dendritic complexity of PCs (Figure 6H and 6I), partially phenocopying the effects of T6B (Figure 1F). Finally, we found that both miR-206 and miR-133 loss-of-function increased CF puncta on PC dendrites (Figure 6J). We observed a similar phenotype for tertiary dendrites when DD-T6B was induced during week 1 (Figure 3F). However, DD-T6B induced during week 2 and week 3 resulted in a decrease in CF puncta (Figures 1H and 3F), suggesting that proper CF innervation requires post-transcriptional mechanisms active during different time windows, and likely involving multiple miRNAs. To our knowledge, this is the first time that a differentially expressed miRNA (miR-206) has been demonstrated to be necessary and sufficient for morphological features that are a hallmark of a specific neuronal subtype (PCs).

## Discussion

The lack of tools for rapidly manipulating global miRNA activity and for accurately mapping miRNA-target networks has prevented systematic investigation of whether miRNAs instruct distinct processes of neuronal differentiation and consequently determine the identity of neuronal subtypes^29^. Here, we overcame these technical hurdles by re-engineering the toolbox to study neuronal miRNAs. Contrary to previous results using kinetically slow approaches^30^, we found that miRNAs are necessary for multiple aspects of PC morphogenesis, suggesting that numerous fundamental roles for miRNAs, especially during postnatal development, remain to be discovered. The ability to inhibit miRNA function rapidly and reversibly using DD-T6B opens new avenues for miRNA research. Not only does DD-T6B offer a powerful way to identify critical windows for miRNA regulation across defined cell types and developmental stages, but it also enables dissection of developmental processes that are overlapping in time.

Mapping miRNA-target networks in defined cell types of the brain illuminates post-transcriptional programs contributing to their development and enables the discovery of the molecular mechanisms instructing their unique morphology and function. Indeed, we have identified a PC-enriched miRNA, miR-206, that is sufficient to increase dendritic complexity in PNs (a PC-like feature) and is necessary for PC-specific morphogenesis, a key determinant of its identity. Remarkably, constitutive KO of miR-29a/b, which is also PC-enriched (3-fold over PNs) but roughly 40-fold more abundant than miR-206 (Figure 6C), causes PC dendritic atrophy at 7 months, but does not affect PC dendritic development^48^. This suggests that miR-206 controls a regulatory network that instructs PC-specific aspects of dendritic morphogenesis. Other notable examples of miRNAs instructing dendritogenesis are miR-9 and miR-137. PN dendritic complexity is reduced by a miR-9 sponge^49^, while it is increased by miR-9 overexpression^79^. Because miR-9 is also enriched in PCs (2.2-fold over PNs; Figure 6C), it might be a contributing factor to their exuberant dendritogenesis. Loss-of-function of miR-137, on the other hand, is known to increase PN dendritic complexity^50,51^. Interestingly, miR-137 is absent in PCs but highly expressed in PNs (Figure 6C), which could be contributing to PN identity by restricting dendritogenesis. These findings demonstrate how differential miRNA expression can shape cell type-specific gene expression to achieve distinctive neuronal morphologies.

A growing body of evidence suggests critical roles for miRNAs in neural progenitor expansion, fate determination, migration, subtype specification, morphogenesis, and circuit formation^4,15,24–29,33–36,49–52,78–90^. However, only a handful of studies show direct evidence supporting the hypothesis that miRNAs instruct neuronal identity. miR-409 mediates the divergence of two cortical neuronal subtypes (callosal projection neurons and corticospinal motor neurons) that are born from the same progenitor but project to either the spinal cord or the contralateral cortex^82,83^. miR-218, which is highly enriched in developing motoneurons^18^, is necessary for establishing motoneuron fate while suppressing interneuron-specific genes^81^. Our data demonstrating that miR-206 is necessary and sufficient to drive the exuberant dendritic morphology characteristic of PCs, is a significant addition to this body of work. Indeed, neuronal morphology, innervation of specific synaptic partners, and subtype-specific molecular profiles are critical determinants of neuronal identity, together with electrophysiological properties, and activity-dependent plasticity^29^.

Finally, it is worth putting our findings in the context of brain evolution. Mammals and soft-bodied cephalopods, two evolutionarily distinct bilaterians with remarkable cognitive functions, expanded their miRNA repertoire very rapidly^7–12^. These novel miRNAs are enriched in neural tissues and during development^12^, leading to the fascinating idea that miRNAs have enabled the emergence of the many cell types that compose complex brains^91^. Consistent with this hypothesis, subsets of miRNAs are developmentally regulated^14–16^ and strongly enriched in different neuronal subtypes^16–18^. Hence, it is possible that evolutionarily novel miRNA sequences and expression patterns contributed to the emergence of new neuronal subtypes^91^. Indeed, the PC-enriched miR-206 is a member of the miR-1/206 seed family that arose through duplication of an ancestral gene common to all bilaterians^6^ and may have been repurposed from the paralogous miR-1 that instructs muscle development^92^ to rewire existing neuronal regulatory networks. Alternatively, PNs may have lost expression of miR-206 (and added others like miR- 137) to produce less branched dendritic arbors that can accommodate considerably different circuit architectures. We believe that the tools developed here will be instrumental for these questions to be explored and in identifying many missing miRNA roles. Furthermore, our tools are not limited to neurons and can be easily repurposed to explore and reexamine the roles of miRNAs in any developmental process and adult function in mice, and hence will have broad impact in many fields.

## Acknowledgements

We are grateful to the past and present members of the Lippi lab for help with plasmid and virus purifications and fruitful discussions. To Dr. Eric Courchesne for advice during the early stages of conceptualization. We thank Dr. Amanda Roberts and the staff at the Mouse Models Core at Scripps Research for performing the behavioral experiments, the Dorris Neuroscience Center vivarium staff for help with mice custom breeding and husbandry, the Salk Institute for Biological Sciences Transgenic Core for generating the Spy3 and cSpy3 mouse lines, Dr. Andrea Ventura at Memorial Sloan Kettering Cancer Center for sharing the Halo-Ago2 mouse line prior to its public availability, and Dr. Eros L. Denchi and Dr. Julia Li for sharing their expertise on CRISPR-Cas9 gene editing and mouse embryonic stem cell culture. We also thank the Genomics Core at Scripps Research for assistance with sequencing the SAPseq libraries and Eclipse Bioinnovations for performing the c-Myc eCLIPseq experiments. We are grateful to Drs Ardem Patapoutian, Lisa Stowers, Xin Jin, and Li Ye for their critical reading of the manuscript. This study was supported by National Institutes of Health grants R01NS121223 and RF1MH126719 (GL), R35GM127090 and R21OD029999 (IJM), F31NS118982 (JXD), by Autism Speaks 12923 (NZ), by the Whitehall Foundation 2018-12-55 (GL).

## Author contributions

Conceptualization: NZ, YX, FZ, IJM, GL. Methodology: NZ, YX, IJM, GL. Investigation: NZ, YX, JXD, MMG, SC, MJJ, ZH, IJM. Analysis: NZ, YX, JXD, MMG, SC, MJJ. Visualization: NZ, YX, JXD, IJM, GL. Funding acquisition: NZ, JXD, IJM, GL. Supervision: IJM, GL. Writing – original draft: NZ, YX, IJM, GL. Writing – review & editing: all authors.

## STAR Methods

### RESOURCE AVAILABILITY

#### Lead contact

Further information and requests for resources and reagents should be directed to and will be fulfilled by the lead contact, Giordano Lippi.

#### Materials availability

All plasmids will be made available at Addgene, and reagents will be available upon request.

#### Data and code availability

SAPseq and CLIPseq data generated as part of this study will be deposited in the Gene Expression Omnibus. CLIPanalyze is available for download at https://bitbucket.org/leslielab/clipanalyze/. Algorithms and key parameters used in this study are available in the method details.

### EXPERIMENTAL MODEL AND SUBJECT DETAILS

#### Animal models

Mice were group-housed on a 12-hr light-dark cycle and fed a standard rodent chow diet. L7-Cre (JAX #010536) and Dicer^fl/fl^ (JAX #006366) were crossed to generate L7^Cre^; Dicer^fl/+^ mice. L7^Cre^; Dicer^fl/+^ and Dicer^fl/fl^ mice were then crossed to generate L7^Cre^; Dicer^fl/fl^ and Dicer^fl/fl^ mice. Homozygous tAgo2 (JAX #017626) transduced with an AAV expressing L7-Cre were used for PC cMyc CLIPseq. VGlut2-ires-Cre (JAX #016963) and tAgo2 (JAX #017626) mice were crossed to generate the VGlut2^Cre^ ^+/-^; tAgo2^+/-^ mice used for PN cMyc CLIPseq. Halo-Ago2 conditional knock-in mice^64^ were obtained from the Andrea Ventura laboratory at MSKCC. VGlut2-ires-Cre (JAX #016963) and Halo-Ago2 mice were crossed to generate VGlut2^Cre+/-^; Ago2^Halo/+^ mice used for Ago2 western blots. Genotyping was performed by Transnetyx using real-time PCR.

The Spy3-Ago2 (constitutive) and cSpy3-Ago2 (conditional) mice were generated via CRISPR/Cas9-mediated gene editing. Initially, several sgRNA candidates were designed via http://crispor.org in the mouse Ago2 gene in intron 1 and exon 2. Through CRISPR screening in E14 mouse embryonic stem cells (mESCs), the sgRNA and repair template with the highest editing efficiency were chosen. For the Spy3-Ago2 line, the sgRNA sequence (5’- GCAAGAACTGGAAAACAGTA-3’) which targets ∼10nt upstream of exon 2 and the repair template with 100 bp homology arms flanking the SpyTag3 knock-in position (Sequence: 5’- TTGGGCCTCGGGATCCAAAGGGCGTGGGAAGGTTAGGGTGCACCATGACTCTCAGGAGCAAGTCTGAGCTTGCACGCCCTGATGCCTGCTGTCTCTCTACTGTTTTCCAGTTCGCGGAGTGCCACATATCGTGATGGTTGACGCTTACAAACGATATAAAGGTAGCGGGGAAAGTGGTAGCGGCGAAAACCTGTATTTTCAGGGCCTTGCTTCTCCTGCTCCGACAACATCACCCATCCCAGGATATGCCTTCAAACCTCCACCTCGGCCGGACTTCGGCACCACCGGGAGAACAATCAAACTAC - 3’) were chosen. The cSpy3-Ago2 line was generated using the conditional by inversion (COIN) approach^69^.For this line, the sgRNA sequence (5’- TTCTTCCAGATCACCAAGAG-3’) targeting intron 1 (∼290 nt upstream of exon 2) and the repair template with the following structure: 5’-(left homology arm)-lox71-(5’ splice site)-(TEV protease cleavage site)-SpyTag3-(3’ splice site)-lox66-(right homology arm)-3’ (Sequence: 5’- ATTGAGCTTGGGGAAGGGAGGCTAGCTGTGACCACAGTGTGAAAGCTCTGAGTCTGGGTGAGAGAGAGGTGAAAGTCCCCAGCAGAGGAATCCTCTCTTGTACCGTTCGTATAGCATACATTATACGAAGTTATTAGCGAAAAAGAAAGAACAATCAAGGGTCCCCAAACTCACCTGCGCCCTGAAAATACAGGTTTTCGCCGCTACCACTTTCCCCGCTACCTTTATATCGTTTGTAAGCGTCAACCATCACGATATGTGGCACTCCGCGACCTGTAGGAAAGAGAAGAAGGCATGAACATGGTTAGCAGAGGGGCCCGTACCGTTCGTATAATGTATGCTATACGAAGTTATGTGATCTGGAAGAACAAGAAGTGTGGGTGGGGGGTTGGCAGAGTCACGGGCCAGCAAGCTGCCTGTCCCTGTCCCTTGTTGGCCCCTGGTAGGTAGGGGT-3’) were chosen (See Figure S4E). The selected sgRNAs were synthesized (Synthego), and mixed 1:1 with Alt-R™ S.p. HiFi Cas9 Nuclease V3 (IDT, 1081060). The repair templates were synthesized via IDT as megamer ssDNA fragments. The sgRNA, Cas9 and repair template were mixed and injected into mouse one-cell staged embryos. The embryos were then transferred into oviducts of pseudo-pregnant mice to produce F0 mice. All microinjection procedures were conducted by The Transgenic Core at Salk Institute for Biological Studies. F0 mice with the strongest knock-in band assessed by genotyping PCR (see below) were selected for backcrossing with C57BL/6J for 10 generations to clean the genetic background.

The constitutive line was genotyped by PCR with the following primers (mAgo2-ex2-F, 5’- GTGGTAAGAAGGTATGAACAGAGG-3’; mAgo2-ex2-R, 5’- TCCAAAGAAATTGGCCTGTAGTTTG-3’) which amplifies a 285-bp band from the WT allele and a 375-bp band from the Spy3-Ago2 allele. The conditional line was genotyped with the following primers (mAgo2-intron1-F, 5’- TGGGACACTGGTTGGAAATG-3’; mAgo2-intron1-R, 5’- CTTACCACCTCATGGCATCC-3’) which amplifies a 459-bp band from the WT allele and a 713-bp band from the cSpy3-Ago2 allele.

VGlut2-ires-Cre and cSpy3-Ago2 mice were crossed to generate VGlut2-Cre; Ago2^cSpy3/+^ mice. Because Ago2 exhibits an imprinted bias of expression from the maternal allele in the mouse brain^66,67^, we used female cSpy3-Ago2 mice for crossing. All experimental protocols were approved by the Scripps Research Institute Institutional Animal Care and Use Committee (IACUC) and were in accordance with the guidelines from the NIH.

#### Cell culture

Cells were maintained in a humidified incubator at 37 °C, 5% CO2.

The wild-type E14 mES cell line was a gift from Dr. Eros Lazzerini Denchi. mESCs were cultured in GMEM (Sigma, G5154) supplemented with 12.5% FBS, Antibiotic/Antimycotic (Gibco, 15240062), GlutaMax (1 mM, Gibco, 35050061), non-essential amino acids (Gibco, 11140050), sodium pyruvate (1mM, Gibco, 11360070), 2-mercaptoethanol (1:100000, Sigma, M7522) and LIF (added fresh 1:1000, Chemicon®, ESG1106). The culture plates were incubated with 0.1% gelatin solution at RT for at least 10 min and the solution was completely removed before plating the cells. The cells were passaged every other day ∼1:10-20.

Primary cortical neuron cultures were prepared from ∼E18 C57BL/6 embryos. The dissection was performed in Leibovitz’s L15 medium (Gibco, 11415064) supplemented with 7mM HEPES (Gibco, 15630080). After media removal, cortices from up to 5 embryos were incubated in 5 mL TrypLE Express (Gibco) for 6 min at 37 °C, mixed by inverting every 1 min. After mechanical dissociation the neurons were plated on 12 mm 1.5 mm thick coverslips (Harvard Apparatus, 64- 0712) coated with poly-D-lysine hydrobromide (0.1 mg/mL, Sigma, P7280) in a 24-well plate. Neurons were cultured at 37 °C in Neurobasal (Gibco, 21103049) medium supplemented with 2% B27 Plus supplement (Gibco, A3582801) and GlutaMax (1 mM, Gibco, 35050061). Half media changes were performed every 3 days. Transfection of neurons for Sholl Analysis was performed with Lipofectamine^TM^ 2000 (Invitrogen, 11668019) at 5 days in vitro (DIV5). Per single well of a 24-well plate, 1 μg DNA was mixed with a 1:50 dilution of Lipofectamine^TM^ 2000 in 100 µl Neurobasal medium and incubated at RT for 30 min. The DNA/Lipofectamine mixture was added into each well containing 400 µl of culture medium and incubated for 1.5 h at 37 °C. For miRNA overexpression experiments, we used QIAGEN miRCURY LNA miRNA mimics (GeneGlobe ID YM00472240 for miR-206 and YM00470608 for miR-133b). The mimics were co-transfected with DNA using the protocol above at a final concentration of 10 nM. All AAV infections were done at DIV5.

## METHOD DETAILS

### Molecular clonings

All PCRs were performed Phusion® High-Fidelity DNA Polymerase (NEB, M0530) and all ligations were performed with T4 DNA ligase (Roche, 10481220001). All plasmids were transformed into One Shot™ Stbl3™ Chemically Competent E. coli (Invitrogen, C737303) and purified using ZymoPURE™ II Plasmid Maxiprep Kit. All plasmid sequences were verified via whole-plasmid sequencing by Primordium Labs.

The sequences for FLAG/HA-T6B-YFP and FLAG/HA-T6Bmut-YFP was obtained here^23^. pAAV[Exp]-SpeI/CBh>XhoI{FLAG/HA-T6B-eYFP}/SalI:WPRE was generated by VectorBuilder. To express T6B in cortical pyramidal neurons we replaced the CBh promoter with the hSyn promoter using SpeI and XhoI. The sequence for hSyn was obtained from the plasmid pLV-hSyn-RFP, a gift from Edward Callaway (Addgene plasmid # 22909). The CBh promoter was replaced with the L7-6 promoter^54^ for PC-specific expression of T6B. The sequence for L7-6 was obtained from pAAV/L7-6-GFP-WPRE (Addgene plasmid #126462) and ordered as a gBLOCK (IDT). T6Bmut was ordered as a gBLOCK (IDT) and cloned into pAAV-L7-6>FLAG/HA-T6B-eYFP:WPRE using NheI and BamHI (two sites flanking T6B).

The inducible T6B (DD-T6B) strategy and design were based on previous studies^61,62^. The DHFR sequence was obtained from the plasmid pAAV-ESARE-DD-EGFP-HA (a gift from Anton Maximov) using PCR. DHFR was cloned into pAAV-hSyn>FLAG/HA-T6B-eYFP:WPRE and pAAV-L7-6>FLAG/HA-T6B-eYFP:WPRE using XhoI and NheI (two sites flanking FLAG/HA).

The mRuby3 construct (pAAV-CAG>mRuby3-V5:WPRE) used to reconstruct PCs for Sholl analysis was a gift from Li Ye. The Cre sequence in pAAV-L7-6>Cre:WPRE was obtained from pENN.AAV.CMVs.Pl.Cre.rBG, a gift from James M. Wilson (Addgene plasmid # 105537) using PCR. Cre was cloned into pAAV[Exp]-L7-6>FLAG/HA-T6B-eYFP:WPRE using XhoI and SalI (two sites flanking FLAG/HA-T6B-eYFP).

In the dual-fluorescence miRNA sensors, TagBFP2 is expressed constitutively from a CMV promoter, while EFS drives expression of mCherry followed by a single miR-124 or miR-206 or miR-133b wildtype or mutated MRE flanked by a 44 bp spacer sequence on the 5’ (AGTGTGGAGGACTTAATCGGTCATTTGCTCACTATGGAATGTGG) and a 33 bp sequence (TTTTTAATCTGTTCCCCTTTGAATACAGAATCC) on the 3’. The spacer sequences were obtained from regions in the Vamp3 gene 3’ UTR that had no annotated MREs. We generated pAAV-rev(CMV>TagBFP2)-EFS>mCherry/BamHI/MluI:WPRE using gBLOCKs. The MREs were ordered as gBLOCKs and with sites for BamHI (5’) and MluI (3’).

The MREs had the following sequences:

<u>miR-124</u> (sequence obtained from Slc32a1 3’ UTR): TCTCTTCTGCTTTGTGCCTTAGT <u>Mutated miR-124:</u> TCTCTgaTGCTTTaTaaCgTgGT (mutated bases in lower case) <u>miR-206</u> (sequence obtained from Tpm4 3’ UTR): TCTCACACTGAAACATTCCACA <u>miR-133b</u> (sequence obtained from Ankrd28 3’ UTR): CATCAAATTGCTGAGGACCAAA

The iRFP670 construct (pAAV-hSyn>iRFP670:WPRE) used to reconstruct PNs for Sholl analysis in miRNA overexpression experiments was obtained by replacing FLAG/HA-T6B-eYFP in pAAV-hSyn>FLAG/HA-T6B-eYFP:WPRE using XhoI and SalI with iRFP670, which was obtained using PCR from a plasmid generated by VectorBuilder.

The sponge design was based on previous studies^93,94^. The sponge sequences were designed using miRNAsong^95^ with the following parameters: bulge from 9-12 nucleotide; 2 miRNA binding sites in a sequence; spacer sequence GAAUAU (see Figure S6E for the sequences). The mutated miR-124 MRE sequence used in the Ctrl sensor was used as a negative control sponge. To achieve directional cloning, we used the compatible ends of the interrupted palindromic SanDI restriction site. We added 5’-GTCCC to the sense and 5’-GACCC to the antisense strand of the miR-124, miR-206 and miR-133 sponge sequences. The strands were ordered as oligonucleotides (IDT) and were resuspended to a final concentration of 100 µM in ultrapure water. The oligonucleotides were mixed at a 1:1 ratio and phosphorylated with T4 PNK (NEB, M0201) and annealed in T4 ligase buffer (NEB, B0202S) in the same reaction with the following parameters: 37 °C for 30 min; 95 °C for 5 min; ramp down to 25 °C at 5 °C min^−1^. For the vector we generated pAAV[FLEX]-CAG>GFP/SanDI:WPRE using gBLOCKs. The vector was prepared for oligonucleotide ligation by digestion with SanDI (KflI, Thermo Scientific^TM^ FD2164) and dephosphorylation with CIP (NEB, M0525). The ligation was performed with a vector/sponge insert ratio of 1:300 at 16 °C for 1h. The ligation reaction was then treated with Plasmid-safe Exonuclease (LGC Biosearch Technologies, E3101K) to digest any residual linearized DNA. After transformation, colony PCR was used to identify the plasmid containing 12 miRNA binding sites.

### AAV Production

All AAVs were produced with the PHP.eB capsid as described^96^. Briefly, AAVs were generated in HEK293FT cells using polyethylenimine transfection. Viral particles were harvested 72 h after transfection from the medium and 120 h after transfection from cells and the medium. Viral particles from the medium were precipitated with 40% polyethylene glycol (Sigma, 89510-1KG-F) in 500 mM NaCl and combined with cell pellets for processing. The cell pellets were suspended in 500 mM NaCl, 40 mM Tris, 2.5 mM MgCl2, pH 8, and 100 U/mL of salt-activated nuclease (Sigma) at 37 °C for 30 min. Afterwards, the cell lysates were clarified by centrifugation at 2,000g and then purified over iodixanol (Optiprep, Sigma; D1556) step gradients (15%, 25%, 40% and 60%). Viruses were concentrated using Amicon filters (EMD, UFC910024) and formulated in sterile phosphate-buffered saline (PBS). Virus titers were measured by determining the number of DNase I–resistant vg using qPCR using a linearized genome plasmid as a standard.

### P0 Intracerebroventricular Injections

PCs can be efficiently targeted by intracerebroventricular (ICV) injections of AAV performed at P0-1. ICV injections were performed as previously described^97^. Injection capillaries (Drummond 3-000-203-g/X) were pulled at the ramp test value on a Sutter P1000 micropipette puller resulting in a tip diameter of ∼90 µm. The injection was performed using a Nanoject III (Drummond). P0 pups were anesthetized by placement on ice fo r 4-5 min until they were immobile and unresponsive to a toe pinch. Then, 1.5 µl of injection mix was injected into each ventricle. Trypan Blue (final concentration of 0.05%) was added to the AAV injection mix to assess ventricle targeting.

### Immunofluorescence

Coverslips with cortical neurons were fixed with prewarmed (37 °C) 4% PFA (1 ml/well) at room temperature for 10 minutes. The coverslips were then washed three times for 5 min each with PBS and mounted onto a slide glass in ProLong^TM^ Diamond Antifade (Thermo Fisher, P36970). Slides were cured overnight at room temperature before imaging.

Mice were transcardially perfused with PBS followed by 4% paraformaldehyde (PFA). Harvested brains were incubated in 4% PFA at 4 °C overnight for post-fixation. Brains were transferred to 30% sucrose in PBS for cryoprotection and left at 4 °C until equilibrated, as assessed by loss of buoyancy. Cryoprotected brains were frozen in a dry ice–hexane bath and sectioned sagittally on a Leica CM1950 at 50 µm. Immunostaining was performed free-floating in 48-well plates. Slices were blocked and permeabilized in 10% Normal Goat Serum (Southern Biotechnologies, 0060-01) and 0.25% Triton-X 100 in PBS. Slices were incubated in primary antibody at 4 °C overnight, washed three times for 5 min each with PBS, incubated with secondary antibody at room temperature for 2 h, washed three times for 5 min each with PBS, and mounted in ProLong^TM^ Diamond Antifade (Thermo Fisher, P36970). Slides were cured overnight at room temperature before imaging. Primary antibodies used were rabbit anti-Eif2c1 (Ago1) (1:200, Proteintech, 19690-1-AP), mouse anti-Ago2 (1:500, Invitrogen, MA5-23515), rabbit anti-GW182 (TNRC6A) (1:500, Novus Biologicals, NBP1-57134), anti-mCherry (1:500, Invitrogen, PA534974), FluoTag®-Q anti-TagFP (1:500, NanoTag Biotechnologies N0401), rabbit anti-calbindin D28K (1:1000, Invitrogen, PA5-85669), guinea pig anti-VGlut2 (1:500, Synaptic Systems, 135404) and rabbit anti-cleaved caspase-3 (1:500, Cell Signaling Technology, 9661). Secondary antibodies used were donkey anti-mouse-AF594 (Invitrogen, A21203), donkey anti-rabbit-AF594 (Invitrogen, A21207), donkey anti-rabbit-AF647 (Invitrogen, A31573) and donkey anti guinea pig-AF647 (Jackson ImmunoResearch, 706605148) all at 1:500. DAPI (final concentration of 1 µM) staining was performed free floating in PBS after the secondary antibody incubation and three washes.

### Confocal microscopy and Image analysis

All images were acquired on a Nikon Instruments A1 confocal laser microscope. Thresholds and laser intensities were established for individual channels and equally applied to entire datasets.

For Sholl analysis of cultured PNs, transfected cells were identified by YFP (for T6B experiments), or iRFP670 (for miRNA overexpression experiments) fluorescence and Z-stacks (at 1 µm steps) were acquired using a 20X dry objective. Sholl analysis was performed on max projections of z-stacks in Fiji using the Sholl plug-in. For PCs, infected cells were identified by mRuby3 and YFP fluorescence (for T6B experiment) or BFP and GFP fluorescence (for sponge experiment) and Z-stacks (at 0.5 µm steps) were acquired using a 60x oil objective. Sholl Analysis was performed on max projections of z-stacks in Fiji using the Sholl plug-in.

Slice area, dendritic height, soma area, lobule IX length, VGlut2 density, VGlut2 height, Calbindin-positive and Caspase-3-positive cells were quantified on the NIS Elements software using the annotations and measurements feature. For VGlut2 density, we manually counted the number of synaptic boutons along the dendrite and reported this number as puncta per dendrite length. The percentage of area of interest covered by VGlut2 synapses was quantified using the Analyze Particles tool in Fiji.

For miRNA sensor experiments virally infected cells were identified by TagBFP2 fluorescence, and Z-stacks (at 1.5 µm steps) were acquired through the entire section using a 20X dry objective. Acquisition settings (laser power, gain, offset) for TagBFP2 and mCherry remained identical for all conditions. Max projections of z-stacks were analyzed in Fiji. ROIs were drawn using the TagBFP2 channel, and the mean fluorescence intensity was measured in both the TagBFP2 and mCherry channels. The ratio of mCherry to TagBFP2 fluorescence was compared between the conditions.

### Behavior

#### Accelerating rotarod test

We used an Accurotar rotarod apparatus (Omnitech Electronics, Inc., Columbus, OH) with a protocol where the rod begins a stationary state and then begins to rotate with a constant acceleration of 10 rpm. When the mice are incapable of staying on the moving rod, they fall onto a teeter-totter arm that moves to break a photobeam. The time of fall (translated to the speed at fall) was recorded by a computer. Mice were tested 6 times per day in two sets of 3: with 1 minute between each test within a set and approximately an hour between each set. This was repeated for two days.

#### Three-chamber sociability test

We used a rectangular, three chambered Plexiglas box, with each chamber measuring 20cmL x 40.5 cmW x 22 cmH. The dividing walls are clear with small semicircular openings (3.5 cm radius) allowing access into each chamber. The middle chamber is empty, and the two outer chambers contain small, round wire cages (Galaxy Cup, Spectrum Diversified Designs, Inc., Streetsboro, OH). The mice were habituated to the entire apparatus for 5 minutes. To assess sociability the mice were returned to the middle chamber, this time with a stranger mouse (C57BL/6J of the same sex being tested, habituated to the wire cage) in one of the wire cages in an outer compartment. Time spent sniffing the stranger mouse and time spent sniffing the empty wire cage was recorded for 5 minutes.

#### Gait Analysis

For gait analysis mice were placed in dish with nontoxic paint so that its paw pads become coated with paint. The mice were then placed at one end of a runway covered in paper and allowed to ambulate until their paws no longer leave marks. From the paint on the paper, stride length (left and right) and stride width (front and back) were measured manually with a ruler.

#### Open field test

For the open field test each mouse was placed in the center of the field of a square white Plexiglas (50 x 50 cm) open field illuminated to 600 lux in the center. Distance traveled, velocity and time spent in center were recorded during a 5-30-minute observation period and analyzed using Noldus Ethovision XT software.

### TMP injections

Trimethoprim lactate salt (AFG Bioscience, 23256-42-0) was reconstituted in PBS prior to each experiment (30 mg/ml) and administered to mice by intraperitoneal injections through a 31 g needle at a dose of 300 μg/gm body weight.

### CLIPseq library preparation

CLIPseq studies were performed by Eclipse Bioinnovations Inc (San Diego, www.eclipsebio.com) according to the published single-end seCLIP protocol^98^ with the following modifications. Approximately 200 mgs of mouse cortical or cerebellum tissue was cryogrinded and UV crosslinked at 400 mJoules/cm2 with 254 nm radiation. Tissue was stored until use at -80 °C. 60 mgs of tissue was aliquoted and lysed using 1mL of eCLIP lysis mix with a modified composition of 6 µl of Proteinase Inhibitor Cocktail and 20 µl of Murine RNase Inhibitor. Samples were then subjected to two rounds of sonication for 4 minutes with 30 second ON/OFF at 75% amplitude. Anti-cMyc Coupled Magnetic beads (ThermoFisher, 88842) or Ago2 antibody (Eclipse Bioinnovations Inc.) pre-coupled to anti-mouse Dynabeads (Thermo Fisher, 11201D) were washed with 500 µl eCLIP Lysis Buffer, added to sonicated lysate, and then incubated overnight at 4 °C. After immunoprecipitation, 2% of the sample was taken as the paired input sample, with the remainder magnetically separated and washed with eCLIP high stringency wash buffers. IP and input samples were cut from the membrane at the relative band size to 75 kDa above. RNA adapter ligation, IP-western, reverse transcription, DNA adapter ligation, and PCR amplification were performed as previously described^98^.

### CLIPseq library processing

The eCLIP cDNA adapter contains a sequence of 10 random nucleotides at the 5’ end. This random sequence serves as a unique molecular identifier (UMI)^99^ after sequencing primers are ligated to the 3’ end of cDNA molecules. Therefore, eCLIP reads begin with the UMI and, in the first step of analysis, UMIs were pruned from read sequences using UMI-tools (v1.1.1)^100^. UMI sequences were saved by incorporating them into the read names in the FASTQ files to be utilized in subsequent analysis steps. Next, 3’ adapters were trimmed from reads using cutadapt (v3.2)^101^ and reads shorter than 18 bp in length were removed. Reads were then mapped to a database of mouse repetitive elements and rRNA sequences compiled from Dfam^102^ and Genbank^103^. All non-repeat mapped reads were mapped to the mouse genome (mm10) using STAR (v2.7.7a)^104^. PCR duplicates were removed using UMI-tools (v1.1.1) by utilizing UMI sequences from the read names and mapping positions. Peaks were identified within eCLIP samples using the peak caller CLIPper (https://github.com/YeoLab/clipper)^105^. For each peak, IP versus input fold enrichment values were calculated as a ratio of counts of reads overlapping the peak region in the IP and the input samples (read counts in each sample were normalized against the total number of reads in the sample after PCR duplicate removal). A p-value was calculated for each peak by the Yates’ Chi-Square test, or Fisher Exact Test if the observed or expected read number was below 5. Peaks were annotated using transcript information from GENCODE Release M25^106^ with the following priority hierarchy to define the final annotation of overlapping features: protein coding transcript (CDS, UTRs, intron), followed by non-coding transcripts (exon, intron).

### Purification of SpyCatcher3 protein

The SpyCatcher3 DNA sequence (5’- GTAACCACCTTATCAGGTTTATCAGGTGAGCAAGGTCCGTCCGGTGATATGACAACT GAAGAAGATAGTGCTACCCATATTAAATTCTCAAAACGTGATGAGGACGGCCGTGAGTTAGCTGGTGCAACTATGGAGTTGCGTGATTCATCTGGTAAAACTATTAGTACATGGATTTCAGATGGACATGTGAAGGATTTCTACCTGTATCCAGGAAAATATACATTTGTCGAAACCGCAGCACCAGACGGTTATGAGGTAGCAACTCCAATTGAATTTACAGTTAATGAGGATGGTCAGGTTACTGTAGATGGCGAAGCAACTGAGGGTGACGCTCATACTGGATCCAGTGGTAGCTAA-3’) was ordered as a gBLOCK (IDT) and cloned into a pET23a-His-TEV plasmid for protein expression. pET23a-His-TEV-SpyCatcher3 plasmid was transformed into Rosetta BL21 E. coli via heat shock at 42 °C for 50 s and cultured on a LB plate with ampicillin (100 µg/ml) and chloramphenicol (25 µg/ml) at 37 °C overnight. The E. coli was then cultured in 30 ml LB with ampicillin and chloramphenicol in a 37 °C shaker (200 rpm) overnight. The next morning, the culture was poured into 3 L LB with ampicillin and chloramphenicol and cultured in a 37 °C shaker until the OD600 reached 0.8 (approximately 4 h). The culture was then supplemented with IPTG (Isopropyl ß-D-1-thiogalactopyranoside, 0.5 mM) and incubated in a 16 °C shaker (200 rpm) overnight.

The E. coli was pelleted at 6000 g for 30 min. The pellet was resuspended in 100 ml lysis buffer (50mM Na_2_HPO4, pH 8.0, 0.3 M NaCl, 5% glycerol, 0.5 mM TCEP, 20 mM imidazole). The resuspended cells were lysed by passing twice through a M-110P lab homogenizer (Microfluidics). The resulting total cell lysate was clarified by centrifugation (30,000 g for 25 min) and the supernatant fraction was applied to a 4.5 mL packed Ni-NTA resin (Qiagen) and gently rocked at 4 °C for 1 h in 50 mL conical tubes. The resin was pelleted by brief centrifugation and the supernatant was discarded. The resin was then washed three times with 50 mL ice-cold Nickel wash buffer (50 mM Tris, pH 8.0, 300 mM NaCl, 20 mM imidazole, 0.5 mM TCEP). Co-purified cellular RNAs were degraded by incubating with 100U micrococcal nuclease (Clontech, 2910A) on-resin in 25 mL of Nickel wash buffer supplemented with 5 mM CaCl_2_ at room temperature for 1 h. The nuclease-treated resin was then washed three times with Nickel wash buffer and then eluted twice sequentially in 8 ml Nickel elution buffer (50 mM Tris, pH 8.0, 300 mM NaCl, 300 mM imidazole, 0.5 mM TCEP). The two elutes were combined with 150 μg TEV protease during an overnight dialysis against 1 L of dialysis buffer (50 mM Tris, pH 8.0, 300 mM NaCl, 20 mM imidazole, 0.5 mM TCEP). On the second day, the dialyzed protein was applied to a 5 ml HisTrap^TM^ column (GE Life Sciences, 17-5248-02), equilibrated with dialysis buffer, and the flow-through containing the SpyCatcher3 protein was collected. This fraction was then further purified by size exclusion chromatography using the HiLoad® 16/600 Superdex® 75 pg column (GE Life Sciences, 28-9893-33) equilibrated with 1x PBS supplemented with 137 mM NaCl, 2.7 mM KCl, 10 mM Na_2_HPO4, 1.8 mM K_2_HPO4. The peak fractions containing SpyCatcher3 were combined and concentrated using a 3 kDa concentrator (Millipore, UFC8003) to ∼8 mg/ml in 1xPBS. The concentrated protein was aliquoted, flash frozen in liquid N2 and stored at –80 °C.

### Western blot

mESCs were harvested and the cell pellets were washed twice with DPBS. The pellets were then lysed in mammalian lysis buffer (50 mM Tris-HCl, pH 7.5, 150 mM NaCl, 1% Triton X-100 and 0.1% Na deoxycholate) containing protease inhibitor cocktail (Thermo Scientific, A32963) at room temperature for 15 min. Lysates were clarified by centrifugation at 12000 g for 10 min. For SpyCatcher3 incubation, 12 µL lysates were mixed with 10 µL of 0.2 mg/ml SpyCatcher3 protein and incubated at room temperature for 1 hour. Cortical tissues were harvested and lysed in mammalian lysis buffer containing protease inhibitor cocktail using a dounce homogenizer.

Lysates were clarified by centrifugation at 12000 g at 4 °C for 15 min. The lysates were further clarified by passing through a 25 mm 0.45 µm syringe filter. All lysates were normalized using bicinchoninic acid (BCA; Pierce BCA Protein Assay Kit) and combined with 4x Laemmli buffer (Bio-Rad). We loaded 50 μg of reduced protein per gel lane and performed transfer with Trans-Blot® Turbo^TM^ Transfer System (Bio-Rad) at 25 V for 7 min. Blocking was performed at room temperature for 60 min with blocking buffer: 5% nonfat dry milk (Bio-Rad) in TBST (150 mM NaCl, 0.1% Tween-20, 50 mM Tris-Cl, pH 7.5). Membranes were then incubated in primary antibody diluted in blocking buffer at 4 °C overnight. After three 5 min washes with TBST, secondary antibodies diluted in blocking buffer were added and incubated for 1 hour at room temperature. After three 5 min washes with TBST, membranes were developed with SuperSignal^TM^ West Pico PLUS Chemiluminescent Substrate (Thermo Scientific, 34579) at room temperature for 5 min. Primary antibodies used were rabbit anti-Ago2 (1:1000, Cell Signaling Technology, 2897S), rabbit anti-beta III Tubulin (1:2000, Abcam, ab18207) as a loading control for cortical tissue; secondary antibody used was goat anti-rabbit-HRP (1:5000, Bio-Rad, 170-6515). Ponceau-S red staining was used for mESCs samples as a loading control.

### Purification of HaloTag-linker-SpyCatcher3 protein

The sequence for HaloTag was amplified from from pENTR4-HaloTag, a gift from Eric Campeau (Addgene: #29644), with the following primers (pET23a-Halo-F, 5’- AATCTCTACTTCCAGGGTACCGGCGCAGAAATCGGTACTGGCTTTC -3’; Halo-linker-R, 5’- GCCGCTACCACTTTCCCCGCTACCGCCGGAAATCTCGAGCGTCG -3’). The SpyCatcher3 sequence was amplified from pET23a-His-TEV-SpyCatcher3 (this paper) with the following primers (linker-SpyCatcher3-F, 5’- GGTAGCGGGGAAAGTGGTAGCGGCGTAACCACCTTATCAGGTTTATC-3’); SpyCatcher3-pET23a-R, 5’- CTCGAATTCGGATCCATCCATGGCTTAGCTACCACTGGATCCAGTATG-3’). These two amplicons were incubated with SfoI digested pET23a-His-TEV plasmid in a HiFi Assembly reaction (NEB, E2621S) to generate a plasmid where HaloTag and SpyCatcher3 is linked with a GSGESGSG linker sequence.

The pET23a-His-TEV-HaloTag-linker-SpyCatcher3 plasmid was transformed into Rosetta BL21 E. coli via heat shock at 42 °C for 50 s and cultured on a LB plate with ampicillin (100 µg/ml) and chloramphenicol (25 µg/ml) at 37 °C overnight. The E. coli was then cultured in 40 ml LB with ampicillin and chloramphenicol in a 37 °C shaker (200 rpm) overnight. The next morning, the culture was poured into 4 L LB with ampicillin and chloramphenicol and cultured in a 37 °C shaker until the OD600 reached 0.6 (approximately 4 h). The culture was then supplemented with IPTG (0.5 mM) and incubated in a 16 °C shaker (200 rpm) overnight.

The E. coli was pelleted at 6000 g for 30 min. The pellet was resuspended in 150 ml lysis buffer (50mM Na_2_HPO4, pH 8.0, 0.3 M NaCl, 5% glycerol, 0.5 mM TCEP, 20 mM imidazole). The resuspended cells were lysed by passing twice through a M-110P lab homogenizer (Microfluidics). The resulting total cell lysate was clarified by centrifugation (30,000 g for 25 min) and the supernatant fraction was applied to a 4 mL packed Ni-NTA resin (Qiagen) and gently rocked at 4 °C for 1 h in 50 mL conical tubes. The resin was pelleted by brief centrifugation and the supernatant was discarded. The resin was then washed three times with 50 mL ice-cold Nickel wash buffer (50 mM Tris, pH 8.0, 300 mM NaCl, 20 mM imidazole, 0.5 mM TCEP). Co-purified cellular RNAs were degraded by incubating with 100U micrococcal nuclease (Clontech, 2910A) on-resin in 25 mL of Nickel wash buffer supplemented with 5 mM CaCl_2_ at room temperature for 1 h. The nuclease-treated resin was then washed three times with Nickel wash buffer and then eluted twice sequentially in 8 ml Nickel elution buffer (50 mM Tris, pH 8.0, 300 mM NaCl, 300 mM imidazole, 0.5 mM TCEP). The two elutes were combined with 150 μg TEV protease during overnight dialysis against 1 L of dialysis buffer (50 mM Tris, pH 8.0, 300 mM NaCl, 20 mM imidazole, 0.5 mM TCEP). On the second day, the dialyzed protein was applied to a 5 ml HisTrap^TM^ column (GE Life Sciences, 17-5248-02), equilibrated with dialysis buffer, and the flow-through containing the SpyCatcher3 protein was collected. This fraction was then further purified by size exclusion chromatography using the Superdex® 200 Increase 10/300 GL column (GE Life Sciences, 28-9909-44) equilibrated with 1x PBS supplemented with 137 mM NaCl, 2.7 mM KCl, 10 mM Na_2_HPO4, 1.8 mM K_2_HPO4. The peak fractions containing HaloTag-linker-SpyCatcher3 were combined and concentrated using a 10 kDa concentrator (Millipore, UFC801024) to ∼15 mg/ml in 1xPBS. The concentrated protein was aliquoted, flash-frozen in liquid N2 and stored at –80 °C.

### Purification of TEV Protease

BL21(DE3) E. coli bearing a His-tagged TEV protease expression plasmid was cultured in 20 ml LB with ampicillin in a 37 °C shaker (200 rpm) overnight. The next morning, the culture was poured into 2 L LB with ampicillin and cultured in a 37 °C shaker until the OD600 reached 0.8 (approximately 4 h). The culture was then supplemented with IPTG (1 mM) and incubated in a 16 °C shaker (200 rpm) overnight.

The E. coli was pelleted at 6000 g for 30 min. The pellet was resuspended in 100 ml lysis buffer (50mM Na_2_HPO4, pH 8.0, 0.3 M NaCl, 5% glycerol, 0.5 mM TCEP, 20 mM imidazole). The resuspended cells were lysed by passing twice through a M-110P lab homogenizer (Microfluidics). The resulting total cell lysate was clarified by centrifugation (30,000 g for 25 min) and the supernatant fraction was applied to a 4 mL packed Ni-NTA resin (Qiagen) and gently rocked at 4 °C for 1.5 h in 50 mL conical tubes. The resin was pelleted by brief centrifugation and the supernatant was discarded. The resin was then washed three times with 50 mL ice-cold Nickel wash buffer (50 mM Tris, pH 8.0, 300 mM NaCl, 20 mM imidazole, 0.5 mM TCEP). The protein was then eluted twice sequentially in 8 ml Nickel elution buffer (50 mM Tris, pH 8.0, 300 mM NaCl, 300 mM imidazole, 0.5 mM TCEP). The protein was further purified by size exclusion chromatography using HiLoad® 16/600 Superdex® 75 pg column (GE Life Sciences, 28-9893-33) equilibrated with 1x Tris buffer (50 mM Tris, pH 8.0, 300 mM NaCl, 0.5 mM TCEP). The peak fractions containing His-TEV-protease were combined and diluted to 0.3 mg/ml with 50% glycerol. The diluted protein was aliquoted, flash-frozen in liquid N2, and stored at –80 °C.

### Preparation of Halo-SpyCatcher3 Resin

400 µL of HaloLink^TM^ resin (Promega, G1912) was washed three times with Halo binding buffer (1x PBS with 0.005% IGEPAL® CA-630 and 0.5mM TCEP). The resin was incubated with 1 mL of 0.5 mg/ml HaloTag-linker-SpyCatcher3 at room temperature for 2 hours, washed five times with Halo binding buffer, and resuspended in 1 mL of Halo binding buffer.

### Halo-SpyCatcher3 Resin Pulldown

Cortical tissues were lysed with mammalian lysis buffer (50 mM Tris-HCl, pH 7.5, 150 mM NaCl, 1% Triton X-100 and 0.1% Na deoxycholate) containing protease inhibitor cocktail (Thermo Scientific A32963) using a dounce homogenizer and incubated for 15 minutes on ice and treated with RQ1 DNase (Promega) for 5 min at 37 °C. The samples were pelleted, and the supernatant was diluted with Halo binding buffer (700 μL per 700 μL lysates). For each sample, 50 µL of Halo-SpyCatcher3 resin was used. The resin was first washed twice with Halo binding buffer. The resin was incubated with the samples for 1 hour at room temperature, washed four times with Halo binding buffer and incubated with 50 µL of TEV (0.3 mg/ml) protease for 1 hour at room temperature to elute Ago2.

### SAPseq library preparation

mESCs were harvested and UV crosslinked twice at 400 mJoules/cm2 with 254 nm radiation in cold PBS on ice. Fresh cortical or cerebellar tissues were harvested, flash frozen in liquid nitrogen, homogenized with a mortar and pestle, and UV crosslinked three times at 400 mJoules/cm2 with 254 nm radiation on dry ice. Cell or tissue pellets were stored at −80 °C until lysis.

Frozen pellets were thawed and lysed in mammalian lysis buffer (50 mM Tris-HCl, pH 7.5, 150 mM NaCl, 1% Triton X-100 and 0.1% Na deoxycholate) containing protease inhibitor cocktail (Thermo Scientific A32963) using a dounce homogenizer and incubated for 15 minutes on ice and treated with RQ1 DNase (Promega) for 5 min at 37 °C. To get the “footprint” of SpyTag3-Ago2, lysates were treated with RNase A (Qiagen 19101) diluted 1:1,000,000 in Halo binding buffer (50 mM Tris-HCl, pH 7.5, 150 mM NaCl, 0.05% IGEPAL® CA-630 and 0.5mM TCEP) for 5 min at 37 °C. The samples were pelleted, and the supernatant was diluted with Halo Binding Buffer (700 μL per 700 μL lysates). The lysates were further clarified by passing through a 25 mm 0.45 µm syringe filter. ∼2% of the lysates were saved for input control library preparation. For each sample, 50 μL Halo-SpyCatcher3 resin was used. The resin was incubated with the diluted lysates at room temperature for 1.5 hours. After incubation, the resin was washed extensively with a series of buffers: Halo binding buffer with 1% Triton X-100 (twice), LiCl wash buffer (100 mM Tris-HCl, pH 8.0, 500 mM LiCl, 1% IGEPAL® CA-630 and 1% Na deoxycholate, three times), 1x PXL buffer (1x PBS with 0.1% SDS, 0.5% Na deoxycholate and 0.5% IGEPAL® CA-630, twice), 5x PXL buffer (5x PBS with 0.1% SDS, 0.5% Na deoxycholate and 0.5% IGEPAL® CA-630, twice) and PNK buffer (50 mM Tris-HCl, pH 7.4, 10 mM MgCl2 and 0.5% IGEPAL® CA-630, twice).

Dephosphorylation was performed on the resin with shrimp alkaline phosphatase (NEB) supplemented with 100 mM CaCl_2_, DNase I (NEB), and murine RNase inhibitor (NEB, M0314) at 37 °C for 20 min. After washes with buffer PNK-EGTA (50 mM Tris-HCl, pH 7.4, 20 mM EGTA and 0.5% IGEPAL® CA-630, twice) and PNK (50 mM Tris-HCl, pH 7.4, 10 mM MgCl_2_ and 0.5% IGEPAL® CA-630, twice), a 3′ RNA adapter with a phosphate on its 5’ (5’- /5Phos/r(N:25252525)r(N:25252525)r(N:25252525)rUrGrGrArArUrUrCrUrCrGrGrGrUrGrCrC rArArGrG/3SpC3/-3’) end was ligated to RNA 3’ ends using T4 RNA ligase 1 (NEB, M0437) at 25 °C for 75 min (with gentle agitation every 10 min). The resin was then sequentially washed with buffer 1x PXL (once), 5x PXL (once), and PNK (three times). RNAs on the resin were treated with T4 PNK (NEB, M0201) supplemented with murine RNase inhibitor at 37 °C for 20 min and washed with Halo binding buffer with 1% Triton X-100 (three times). To release the Ago-miRNA-mRNA complexes, the resin was incubated in TEV elution buffer (Halo binding buffer, 1% Triton X-100, murine RNase inhibitor, 1.5 µg TEV protease) at room temperature for 1 hour (with gentle agitation every 10 min). The elute and input samples were combined with 4x Laemmli buffer (Bio-Rad) and run on an SDS-PAGE gel. For each sample, the gel was cut between 70 kDa and 150 kDa and crushed with a P1000 tip. The gel fragments were incubated with 4 mg/mL proteinase K (NEB, P8107) in PK buffer (100 mM Tris-HCl, pH 7.5, 50 mM NaCl and 10 mM EDTA) at 37 °C for 20 min and further inactivated by 7 M urea dissolved in PK buffer at 37 °C for 20 min. Free RNAs were extracted using phenol/chloroform and purified with the RNA Clean & Concentrator-5 kit (Zymo Research, R1015).

To prepare the input control library, the purified RNA was dephosphorylated with shrimp alkaline phosphatase (NEB) supplemented with 100 mM CaCl_2_, DNase I and murine RNase inhibitor at 37 °C for 20 min and cleaned up using the MyONE^TM^ Silane beads (Invitrogen, 37002D) as described^107^. Then, the 3′ RNA adapter was ligated to the purified RNAs using T4 RNA ligase 1 (NEB, M0437) at 25 °C for 75 min (with gentle agitation every 10 min). The ligated RNAs were purified using the MyONE^TM^ Silane beads. Then, the RNAs were phosphorylated with T4 PNK (NEB, M0201) supplemented with murine RNase inhibitor at 37 °C for 20 min and purified using MyONE^TM^ Silane beads.

A 5′ RNA adapter (5’-/5biotin/rGrUrUrCrArGrArGrUrUrCrUrArCrArGrUrCrCrGrArCrGrArUrCrNrNrNrNrNrNrNrN rNrN-3’) was ligated to the purified elute and input RNAs using T4 RNA ligase 1 at 16 °C overnight. The next day, the RNAs were purified with the RNA Clean & Concentrator-5 kit. The purified RNAs were reverse transcribed using the DP3 primer (5’- CCTTGGCACCCGAGAATTCCA-3’, 0.5 μM) and Superscript III reverse transcriptase (Invitrogen, 18080-044) and treated with Exonuclease I (NEB, M0293) at 37 °C for 30 min and purified using MyONE^TM^ Silane beads. The resulting cDNAs were amplified with primers DP3 and DP5 (5’-GTTCAGAGTTCTACAGTCCGACGATC-3’, 0.5 μM) and KAPA Hifi HotStart DNA polymerase (Roche, KK2601) to the optimal amplification point. The optimal amplification cycle (defined as the cycle before the PCR reaction reaching a plateau) was preliminarily determined by a diagnostic PCR visualized on a 10% TBE polyacrylamide gel (Invitrogen, EC6275BOX). PCR products of miRNAs (expected size range: 75-81 bp) and targets (expected size range: 88∼135 bp) from the elute samples and inputs (expected size range: 88∼135 bp) were resolved on a 10% TBE polyacrylamide gel and extracted separately using the DNA Clean & Concentrator-5 kit (Zymo Research).

To construct the libraries for high-throughput sequencing, DNA primers TS Primer-X (“X” stands for barcode index; CAAGCAGAAGACGGCATACGAGATNNNNNNGTGACTGGAGTTCCTTGGCACCCGAG AATTCCA, N=standard Illumina barcodes) and TS Primer-1 (AATGATACGGCGACCACCGAGATCTACACGTTCAGAGTTCTACAGTCCGACGATC) containing Illumina barcodes, sequencing primer binding sites and Illumina TruSeq indexes for multiplexing were introduced to the SAPseq miRNA and target libraries by PCR with the KAPA Hifi HotStart DNA polymerase (Roche KK2601). PCR products were purified using AmpureXP beads (Beckman A63880). The sequencing libraries were then run on a 10% TBE polyacrylamide gel and purified using the DNA Clean & Concentrator-5 kit (Zymo Research D4013).

SAPseq target and miRNA libraries, along with the matched input control libraries, were submitted to the Genomics Core at Scripps Research for high-throughput sequencing. After quantification and quality control, libraries were pooled and run on a NextSeq 2000 in a 100 bp single end run.

### SAPseq miRNA library processing

#### UMI extraction

The first 8 nt that encodes the UMI in each read was extracted by UMI-tools^100^ and appended to the original read name, which was used to distinguish duplicated reads produced at PCR amplification steps.

#### Adapter removal and read quality control

The 3′ adapter and bases with Phred quality score lower than 20 were trimmed from reads using cutadapt v2.10^101^. Reads less than 15bp were discarded.

#### miRNA abundance estimates

Processed small RNA reads were aligned to a miRNA genome index built from 1,915 murine pre-miRNA sequences from miRbase version 22 (ftp://mirbase.org/pub/mirbase/21/)^108^ using Bowtie v2.4.2^109^. The miRNA counts were normalized to counts per million (cpm).

miRNA seed family data were downloaded from the TargetScan website (mmu80)^110^. For miRNA family level analysis in Figure 4C, read counts mapping to members of the same miRNA family were summed up.

### SAPseq target library processing

Reads in the target libraries were processed and filtered following the “UMI extraction” and “Adapter removal and read quality control” steps described in the “SAPseq miRNA library processing” section.

#### Alignment

Processed reads were aligned to the UCSC mm10 mouse genome using STAR v2.5.2a^104^.

#### PCR duplicate removal

Reads mapped to the same locus with identical UMIs were considered PCR duplicates and therefore collapsed using the UMI-tools dedup function with default 1-mismatch allowed in the UMI region for error correction. Representative reads of these unique events were written into a new BAM file, which was used for peak calling.

#### Peak calling

Peaks were called using the unpublished package CLIPanalyze (https://bitbucket.org/leslielab/clipanalyze). The function findPeaks () was run with the following parameters: bandwidth = 30, count.exons.only = FALSE, nthreads = 4, min.width = 15, merge.dist = 20. Differential read count analysis was performed between target and input control read counts in putative peaks using DESeq2 v1.36.0^111^, and FDR-corrected p-values (adjusted p-values) were assigned to each peak. Non-miRNA peaks of read count log2FC > 0 and adjusted p-value < 0.1 in target versus input control were selected for downstream analysis. Peaks were annotated as overlapping with 3′UTR, 5′UTR, exons, introns and intergenic, in that order, using GENCODE vM10 annotation^106^. Peaks annotated as exons were further separated as protein-coding (CDS) and noncoding-RNA (ncRNA). Finally, the peaks were annotated with a list of potential miRNA family seed matches for miRNA families with cpm>100.

#### SAPseq coverage analysis

bigWig files for visualization of target and input control libraries were produced using deepTools bamCoverage v3.1.3^112^ with parameter “--binSize 1 --normalizeUsing CPM”. SAPseq libraries were visualized using IGV^113^.

### SAPscore calculation

SAPscores were calculated for each replicate individually. Briefly, we first calculated a SAP RPKM for each gene, and then calculated the residual for each gene after regressing SAP RPKM on gene expression, as measured by RNAseq TPM.

SAPseq RPKMs were calculated by adding all counts falling on peaks in the 3’ UTR of each gene called with >2 log2 fold change over input and <0.05 adjusted p-value, normalizing by the 3’ UTR length in kilobases and each replicate’s library size, and then multiplying by a 10^6^ scaling factor, as follows.

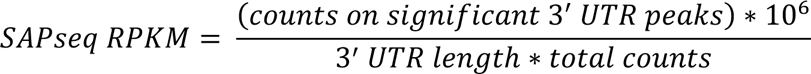

PC P14 RNA sequencing raw counts^75^ were converted into TPMs by normalizing for transcript length and library size with a per million scaling factor. PN P15 RNA sequencing RPKM was obtained here^114^. We then used Deming regression from the mcr v1.2.2 R package to fit a linear regression of SAPseq RPKM on RNAseq TPM and defined the SAPscore as the optimized residual of each gene’s RPKM from this regression. The mean SAPscore across replicates were used for downstream analysis.

### Gene Ontology (GO) Term Analysis

GO term analysis was performed using the enrichGO and simplify functions from ClusterProfiler v3.18.1. Each analysis was performed using the Biological Process ontologies and the Benjamin-Hochberg false discovery rate p-value adjustment with p-value cutoff of 0.01; the simplify function was used with standard settings and cutoff of 0.7 to remove redundant GO terms. For PC gene lists, the universe was set as all genes detected in RNAseq^75^. For PN gene lists, the universe was set as all genes detected in RNAseq from Fertuzinhos et al. ^114^.

### Bipartite network construction

The bipartite network was generated using the top 30 miRNAs and targets whose 3’ UTR peaks have predicted MREs for those miRNAs in P15 PCs using the R bipartite package (v2.17)^115^.

### Differential expression analysis of miRNAs

To analyze differential expression of miRNAs, we used DESeq2 (v1.30.1)^111^ and input the raw miRNA counts from each replicate. MiRNAs with adjusted p-value < 0.05 were considered as significantly differentially expressed.

### Statistical analysis

Statistical analyses were performed in GraphPad Prism and R. All datasets were tested for outliers, normal distribution, and equal variance. For Sholl analysis we performed a mixed model effects analysis followed by Šídák’s multiple comparisons test if there was a significant condition x distance from soma interaction. All statistical and quantitative analyses are described in detail at the end of each figure legend.

