## Supplemental Figures for "MicroRNAs are necessary for the emergence of Purkinje cell identity"

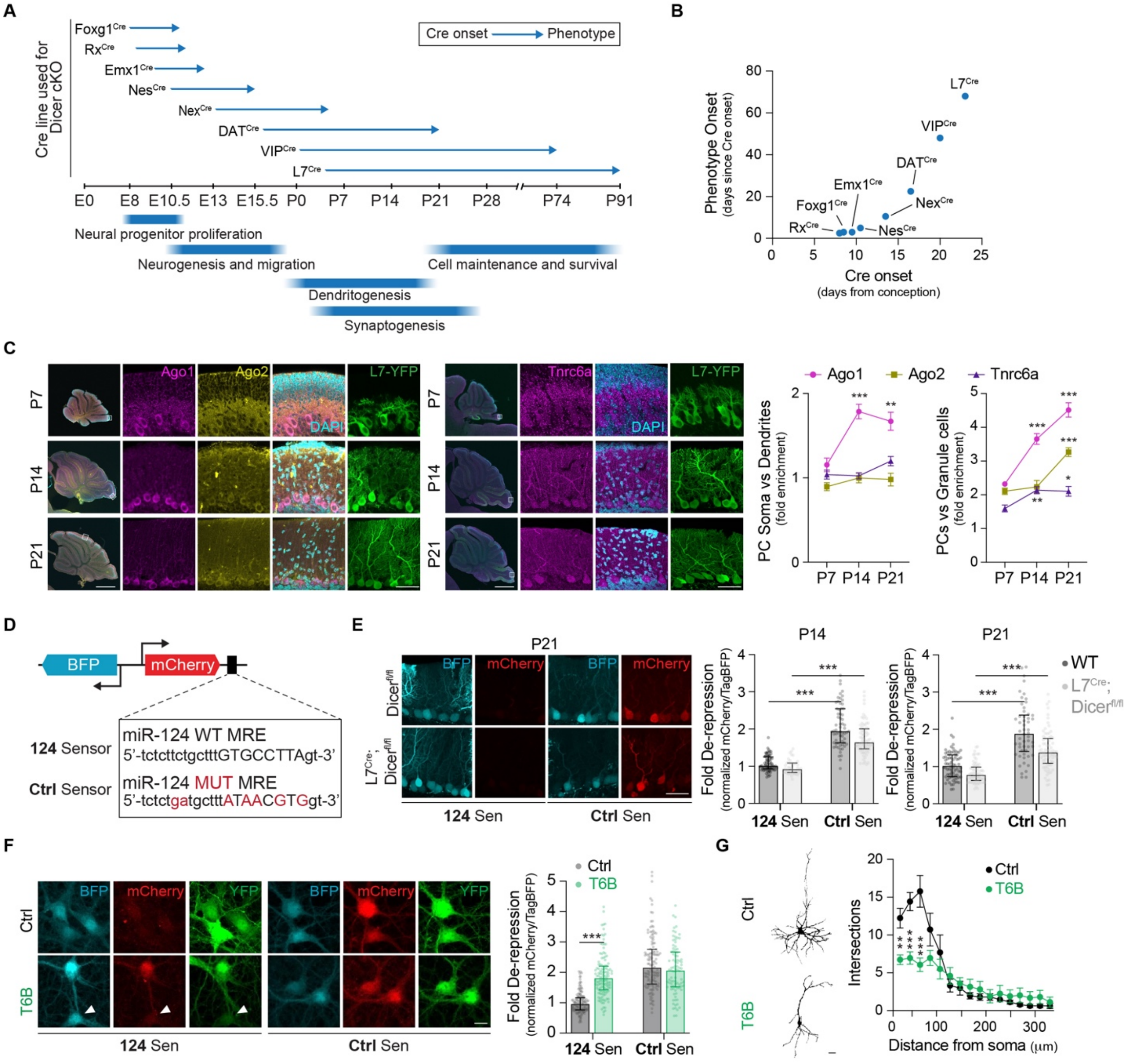


**Figure S1**. **The miRNA machinery is intact and functional in developing PCs**, **related to Figure 1.**

(A) Timeline of neuronal differentiation events in the mouse with examples of Dicer cKO studies[25–28,30,32,34,35]. For each Cre line used, the phenotype onset is shown. Only Dicer cKO with Cre lines that turn on embryonically affects dendritic and synaptic morphogenesis, two critical aspects of neuronal identity.

(B) The phenotype onset gets progressively more delayed with Cre lines that turn on later in the developmental timeline.

(C) Immunostaining and quantification of Ago1, Ago2 and Tnrc6a in P7, P14 and P21 PCs. The miRISC complex is enriched in PCs. Scale bar, 1 mm for whole slice images; 50 μm for PCs close-up. N=12 cells.

(D) Schematic of the dual-fluorescence miRNA sensor (top). BFP functions as a transduction reporter and is not regulated by miRNAs. mCherry harbors a miR-124 MRE and functions as a miR-124 sensor. Sequences of the wild-type (WT) and mutated (MUT, mutated bases in red) miR-124 MREs (bottom) cloned in the 124 and Ctrl sensors.
(E) Quantification of the miR-124 sensor activity in P14 and P21 $Dicer^{fl/fl}$ (WT) and $L7^{Cre}$; $Dicer^{fl/fl}$ PCs shows intact miRNA function. Scale bar, 50 µm. N=45-80 cells from 2 mice.
(F) Functional validation of T6B in cultured cortical PNs using the miR-124 sensor at DIV11. T6B de-repressed mCherry translation. mCherry remains repressed in neurons not expressing T6B (white arrow). Scale bar, 10 µm. N=91-145 cells.
(G) Representative images and quantification of dendritic complexity (Sholl analysis) in cultured cortical PNs at DIV11. T6B decreased dendritic complexity. Scale bar, 10 µm. N=8 cells.
Data are mean ± SEM for (C) and (G); median with interquartile range for (E) and (F). Statistics for (C): Welch's t-test, (E) and (F): Mann-Whitney test; (G): Mixed-effects model with Šídák's multiple comparisons test. $^{**}p \leq 0.01$; $^{***}p \leq 0.001$.

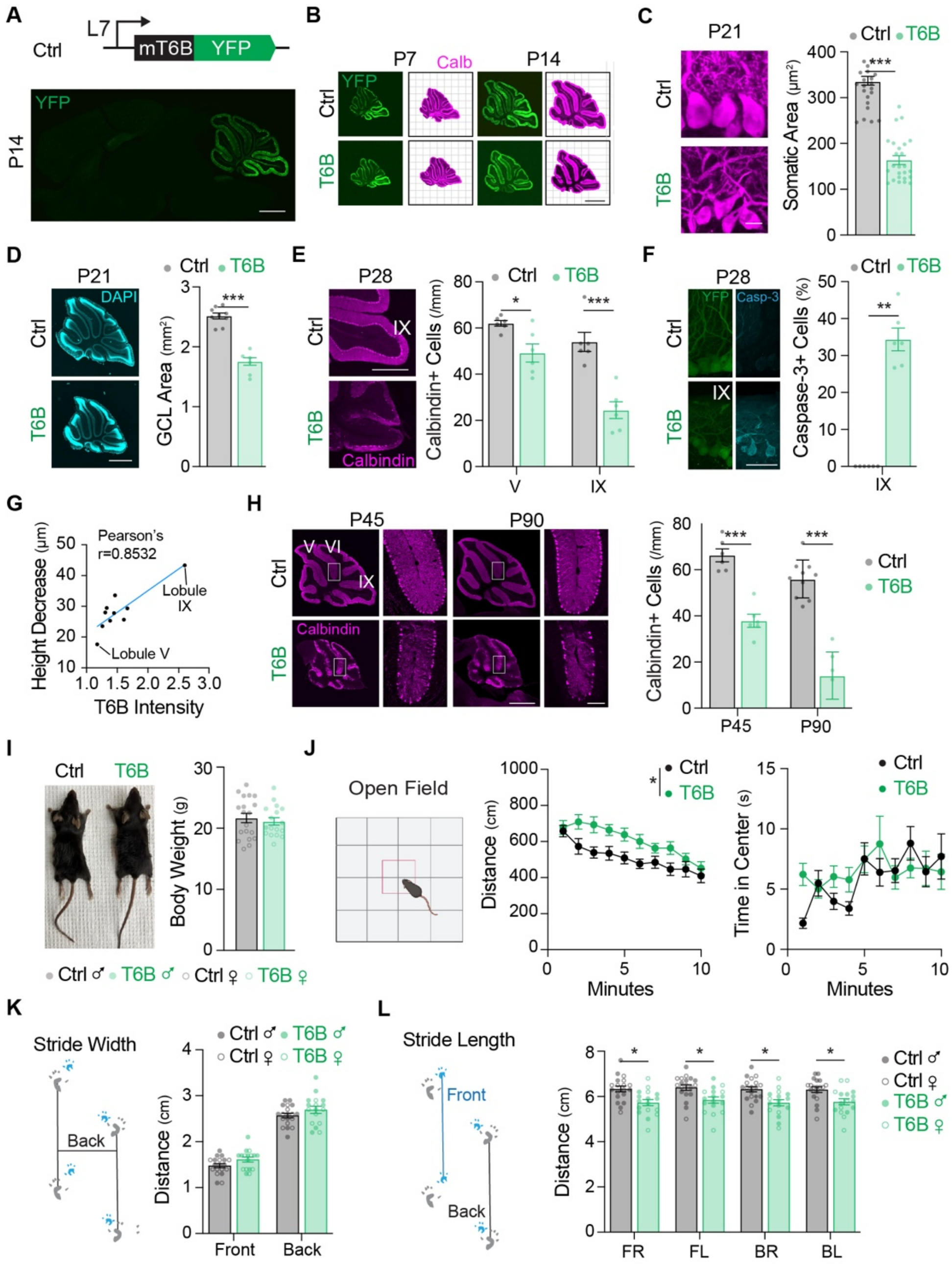

**Figure S2. T6B induces developmental phenotypes in PCs and defects in behaviors linked to PC function, related to Figure 1.**
(A) Whole brain sagittal slice showing expression of Ctrl (L7-mT6B) exclusively in PCs. Scale bar, 1 mm.
(B) Calbindin-stained sagittal sections of Ctrl- and T6B-transduced mice collected at P7 and P14. T6B induced a reduction in cerebellar area (quantification in Figure 1D). Scale bar, 1 mm.
(C) Calbindin-stained sagittal sections showing smaller PC somas in T6B-transduced mice at P21. Scale bar, 10 µm. N=20 cells from 3 mice.
(D) DAPI-stained sagittal sections showing a reduction in the granule cell layer (GCL) area in T6B-transduced mice at P21. Scale bar, 1 mm. N=9 sections from 3 mice.

(E) Calbindin-stained sagittal sections at P28. Quantification of PCs in lobule V and in lobule IX shows that T6B causes PC loss. Scale bar, 500 µm. N=6 sections from 3 mice.
(F) Cleaved caspase-3 staining in lobule IX shows that T6B induced apoptosis. Scale bar, 50 µm. N=6 sections from 3 mice.
(G) Dose-dependency of the T6B-induced phenotype. T6B transduction is most efficient in lobule IX and least efficient in lobule V. The decrease in dendritic height positively correlates with T6B expression levels.
(H) Calbindin-stained sagittal sections showing PC loss at P45 and P90. Scale bar, 1 mm.
(I) T6B-transduced mice did not show changes in body weight at P64-70. N=18 mice.
(J) Open field test at P60. T6B-transduced mice were moderately hyperactive (middle) but spent similar amount of time in the center of the open field to controls, a sign that T6B did not induce anxiety-like behaviors (right). N=18 mice
(K) T6B-transduced mice did not show changes in stride width at P60. Left, schematic of how measurements are taken from paw prints. N=18 mice.
(L) T6B-transduced mice had shorter stride length for all four paws. F: front, B: back, R: right, L: left. N=18 mice.
Data are mean ± SEM. Statistics for (C), (D), (E) and (H): Welch's t-test; (G): Pearson's correlation coefficient; (J) and (L): Two-way ANOVA. $^{*}p \leq 0.05$; $^{**}p \leq 0.01$; $^{***}p \leq 0.001$.

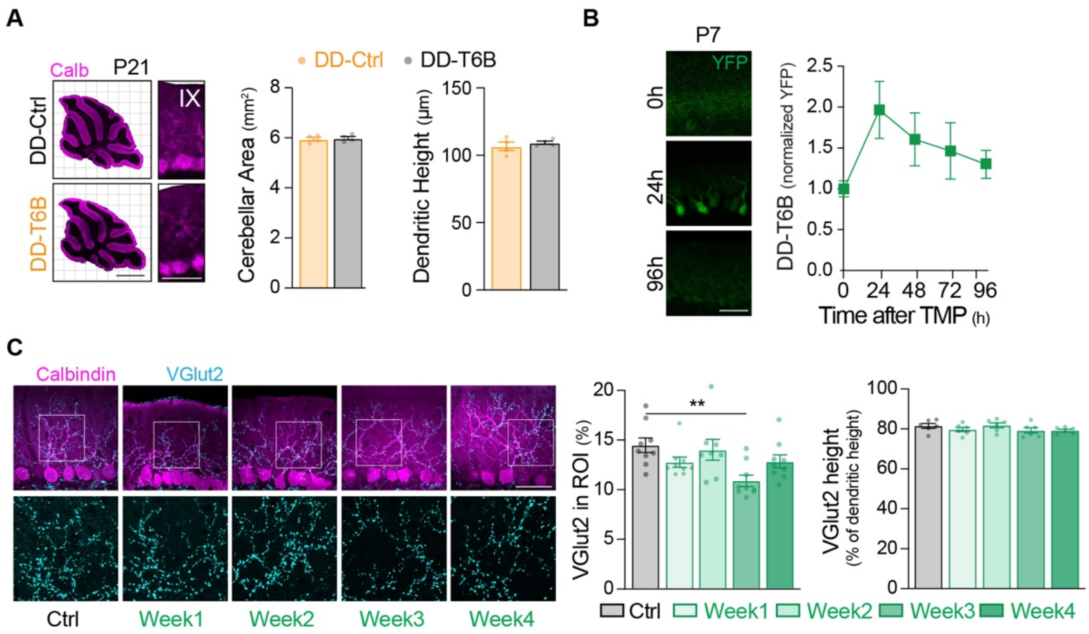


**Figure S3. Inducible T6B allows temporally restricted miRNA loss-of-function, related to Figures 2 and 3.**

(A) Calbindin-stained sagittal sections of DD-Ctrl- and DD-T6B-transduced mice collected at P21. Without TMP, DD-T6B had no effect on cerebellar area or dendritic height. Scale bar, 1mm for area; 50 µm for dendritic height. N=4 sections from 2 mice.

(B) DD-T6B induction time course in PCs at P7. Scale bar, 50 µm. N=35-48 cells.

(C) Calbindin-stained sagittal slices at P28 show that DD-T6B induction during week 3 reduces the percentage of CF VGlut2 puncta. CFs reach the same percentage of the PC dendrites at each week. Scale bar, 50 µm. N=9 sections from 3 mice.

Data are mean ± SEM. Statistics: Two-way ANOVA. $^*p \leq 0.05$; $^{**}p \leq 0.01$; $^{***}p \leq 0.001$.

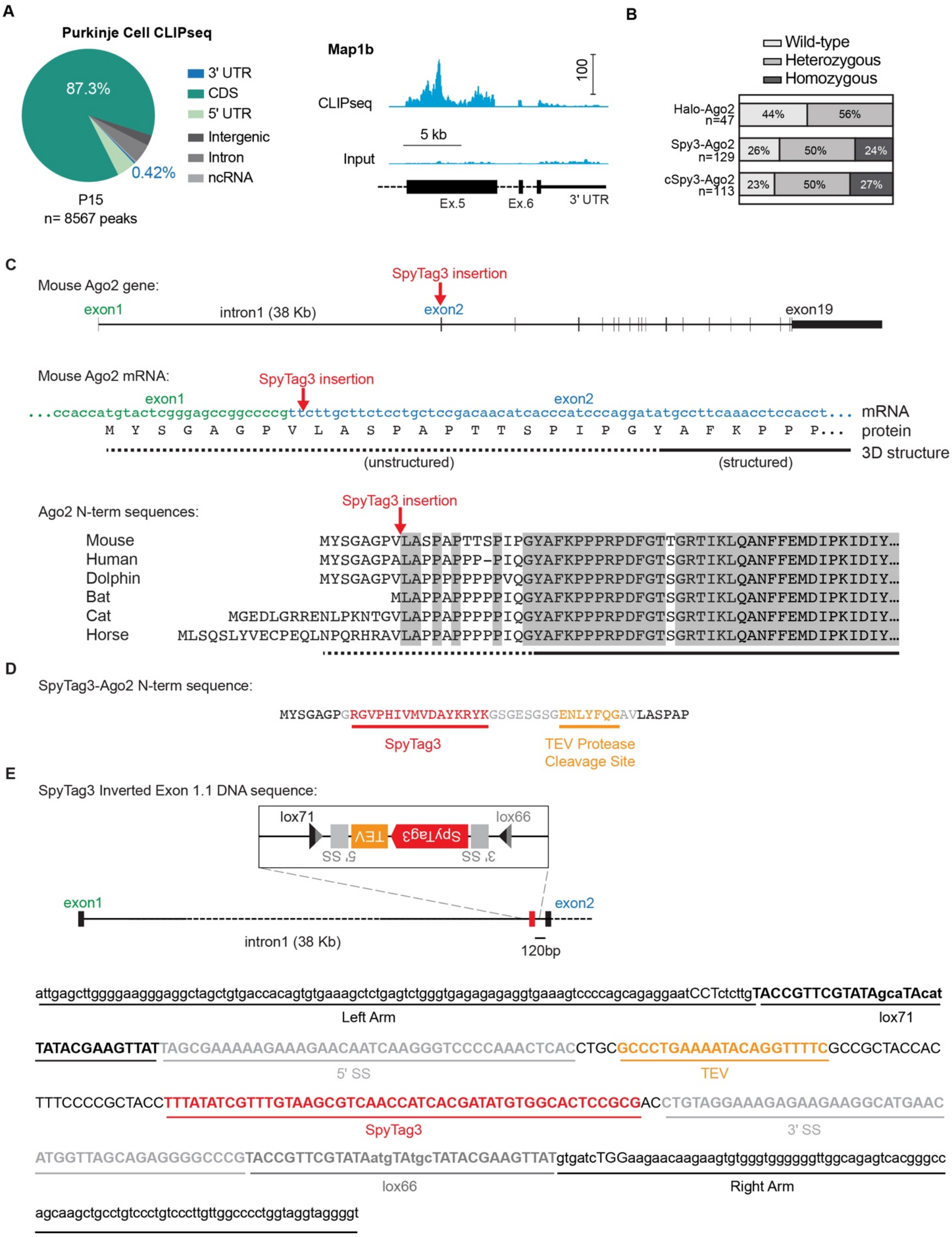

A
Purkinje Cell CLIPseq
87.3%
0.42%
3' UTR
CDS
5' UTR
Intergenic
Intron
ncRNA
P15
n= 8567 peaks
Map1b
CLIPseq
100
5 kb
Input
Ex.5
Ex.6
3' UTR
B
Wild-type
Heterozygous
Homozygous
Halo-Ago2 n=47
44%
56%
Spy3-Ago2 n=129
26%
50%
24%
cSpy3-Ago2 n=113
23%
50%
27%
C
Mouse Ago2 gene:
SpyTag3 insertion
exon1
intron1 (38 Kb)
exon2
exon19
Mouse Ago2 mRNA:
SpyTag3 insertion
exon1
exon2
...ccaccatgtactcgggagccggccccgttcttgcttctcctgctccgacaacatcacccatcccaggatatgccttcaaacctccacct... mRNA
M Y S G A G P V L A S P A P T T S P I P G Y A F K P P P... protein
3D structure
(unstructured)
(structured)
Ago2 N-term sequences:
SpyTag3 insertion
Mouse MYSGAGPVLASPAPTTSPIPGYAFKPPPRPDFGTTGRTIKLQANFFEMDIPKIDIY...
Human MYSGAGPALAPPAPPP-PIQGYAFKPPPRPDFGTSGRTIKLQANFFEMDIPKIDIY...
Dolphin MYSGAGPVLAPPPPPPPVQGYAFKPPPRPDFGTSGRTIKLQANFFEMDIPKIDIY...
Bat MLAPPAPPPPPIQGYAFKPPPRPDFGTSGRTIKLQANFFEMDIPKIDIY...
Cat MGEDLGRRENLPKNTGVLAPPAPPPPPIQGYAFKPPPRPDFGTSGRTIKLQANFFEMDIPKIDIY...
Horse MLSQSLYVECPEQLNPQRHRAVLAPPAPPPPPIQGYAFKPPPRPDFGTSGRTIKLQANFFEMDIPKIDIY...
D
SpyTag3-Ago2 N-term sequence:
MYSGAGPGRGVPHIVMVDAYKRYKGSGESGSGENLYFQGAVLASPAP
SpyTag3
TEV Protease Cleavage Site
E
SpyTag3 Inverted Exon 1.1 DNA sequence:
lox71
lox66
TEV
SpyTag3
5' SS
3' SS
exon1
exon2
intron1 (38 Kb)
120bp
attgagcttggggaagggaggctagctgtgaccacagtgtgaaagctctgagtctgggtgagagagaggtgaaagtccccagcagaggaatCCTctcttgTACCGTTCGTATAgcaTAcat
Left Arm
lox71
TATACGAAGTTATTAGCGAAAAAGAAAGAACAATCAAGGGTCCCCAAACTCACCTGCGCCCTGAAAATACAGGTTTTCGCCGCTACCAC
5' SS
TEV
TTTCCCCGCTACCTTTATATCGTTTGTAAGCGTCAACCATCACGATATGTGGCACTCCGCGACCTGTAGGAAAGAGAAGAAGGCATGAAC
SpyTag3
3' SS
ATGGTTAGCAGAGGGGCCCGTACCGTTCGTATAatgTAtgcTATACGAAGTTATgtgatcTGGaagaacaagaagtgtgggtgggggttggcagagtcacgggcc
lox66
Right Arm
agcaagctgcctgtccctgtcccttgttggcccctggtaggtaggggt

**Figure S4. Design of the Spy3-Ago2 and cSpy3-Ago2 mouse lines, related to Figure 4.**
(A) Frequency across genomic annotations of the peaks mapped via Ago2 CLIPseq from P15 tAgo2$^{+/+}$ mice transduced with L7-Cre AAV. 3' UTR, 3' untranslated region; 5' UTR, 5' untranslated region; CDS, coding sequence; ncRNA, non-coding RNA. The majority of peaks mapped to CDS while <1% mapped to the 3' UTR. Example on the right: Genome browser view of Map1b shows peaks in the CDS but not in the 3' UTR.
(B) Frequency of genotypes from heterozygous intercrosses of Halo-Ago2, Spy3-Ago2, and cSpy3-Ago2 mice.
(C) The SpyTag3 is inserted at the beginning of the exon 2 to avoid interfering with the Ago2 promoter. This area was chosen because it is unstructured and poorly conserved, suggesting non-essential functions. The C-terminus of Ago2 is not suitable for tagging because any additional peptide would likely affect the miRNA binding pocket.
(D) Amino acid sequence of the SpyTag3 insert.
(E) DNA sequence of the construct used to generate the cSpy3-Ago2 mouse line. 3' SS, 3' splice site; 5' SS, 5' splice site; TEV, TEV protease cleavage site.

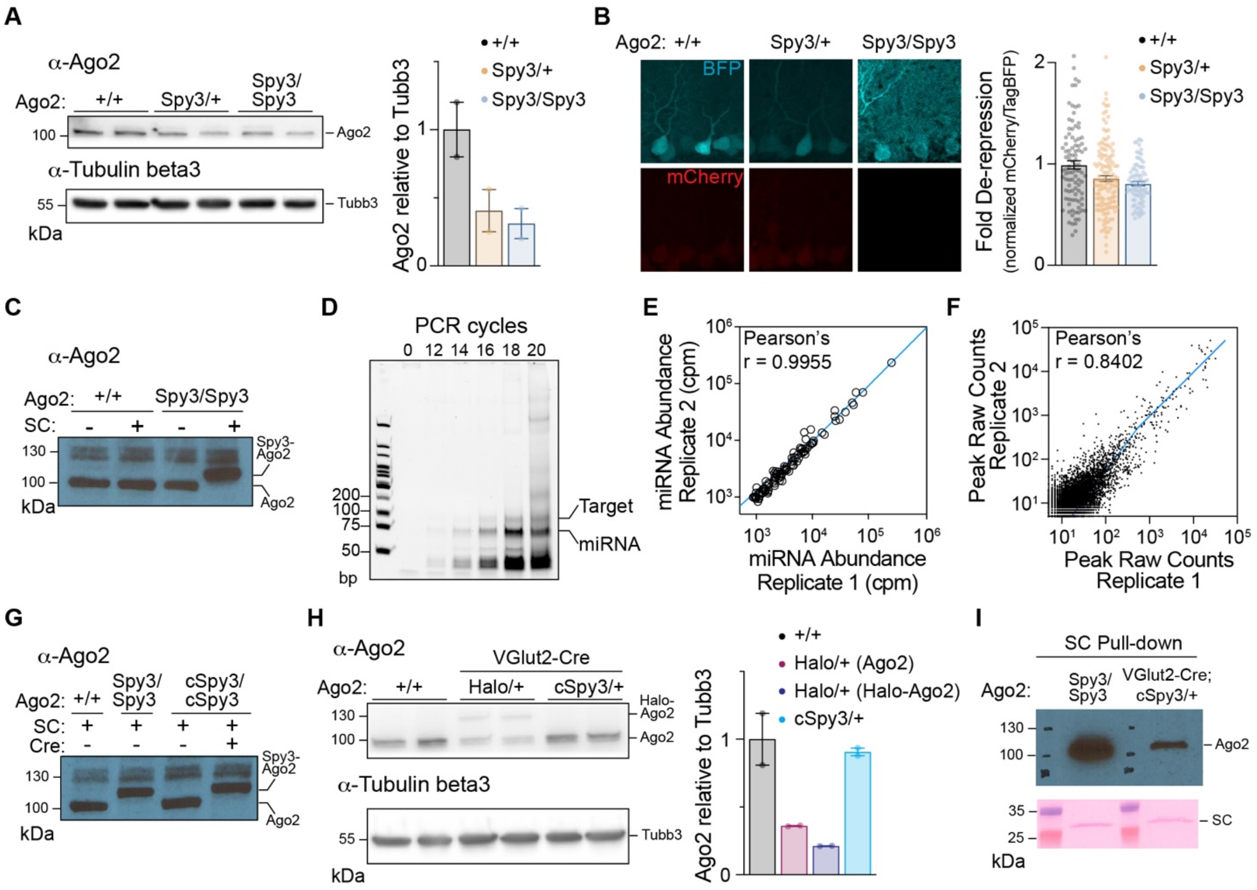


**Figure S5. Validation of the Spy3-Ago2 and cSpy3-Ago2 mESCs and mouse lines, related to Figure 4.**

(A) α-Ago2 Western blot (with quantification) of cortical lysates from Ago2$^{+/+}$ (wild-type), Ago2$^{Spy3/+}$ (heterozygous constitutive Spy3-Ago2) and Ago2$^{Spy3/Spy3}$ (homozygous constitutive Spy3-Ago2) mice at P14. Endogenous Ago2 levels are lower in Ago2$^{Spy3/+}$ and Ago2$^{Spy3/Spy3}$ mice.

(B) miR-124 sensor activity in Ago2$^{+/+}$, Ago2$^{Spy3/+}$, and Ago2$^{Spy3/Spy3}$ PCs. miRNA activity is similar in all three mouse lines. Data are mean ± SEM. N=84-139 cells from 2 mice.

(C) α-Ago2 Western blot of Ago2$^{+/+}$, Ago2$^{Spy3/Spy3}$ E14 mESC lysate treated with or without SpyCatcher3 protein (SC). Because Spy3 is small, binding to SC is necessary to see the Ago2 band shift.

(D) 10% denaturing PAGE showing SAPseq libraries after various rounds of PCR amplification. Bands corresponding to miRNA and miRNA-target libraries indicated.

(E) Pairwise comparison of miRNA abundance quantified from two independent miRNA libraries generated from mESCs show positive correlation.

(F) Pairwise comparison of peaks called in two independent target libraries generated from mESCs show strong positive correlation.

(G) α-Ago2 Western blot of SC-treated lysate from E14 Ago2$^{cSpy3/cSpy3}$ mESCs cells after transfection and selection with Cre recombinase. SC-treated lysates from E14 Ago2$^{+/+}$ and Ago2$^{Spy3/Spy3}$ mESCs included as controls.

(H) α-Ago2 Western blot (with quantification) of cortical lysates from $Ago2^{+/+}$, $VGlut2^{Cre}$ ; $Ago2^{Halo/+}$, and $VGlut2^{Cre}$; $Ago2^{cSpy3/+}$ mice at P14. Ago2 levels are lower in $VGlut2^{Cre+/-}$; $Ago2^{Halo/+}$ but normal in $VGlut2^{Cre+/-}$; $Ago2^{cSpy3/+}$ mice.

(I) α-Ago2 Western blot of SpyCatcher3 pulldown from cortical lysates of $Ago2^{Spy3/Spy3}$ and $VGlut2^{Cre+/-}$; $Ago2^{cSpy3/+}$ mice.

Statistics for (E) and (F): Pearson's correlation coefficient. The blue line represents best-fit linear regression; (I) and (J): Spearman's rank correlation coefficient. The blue line represents best-fit linear regression, with 95% confidence interval in gray.

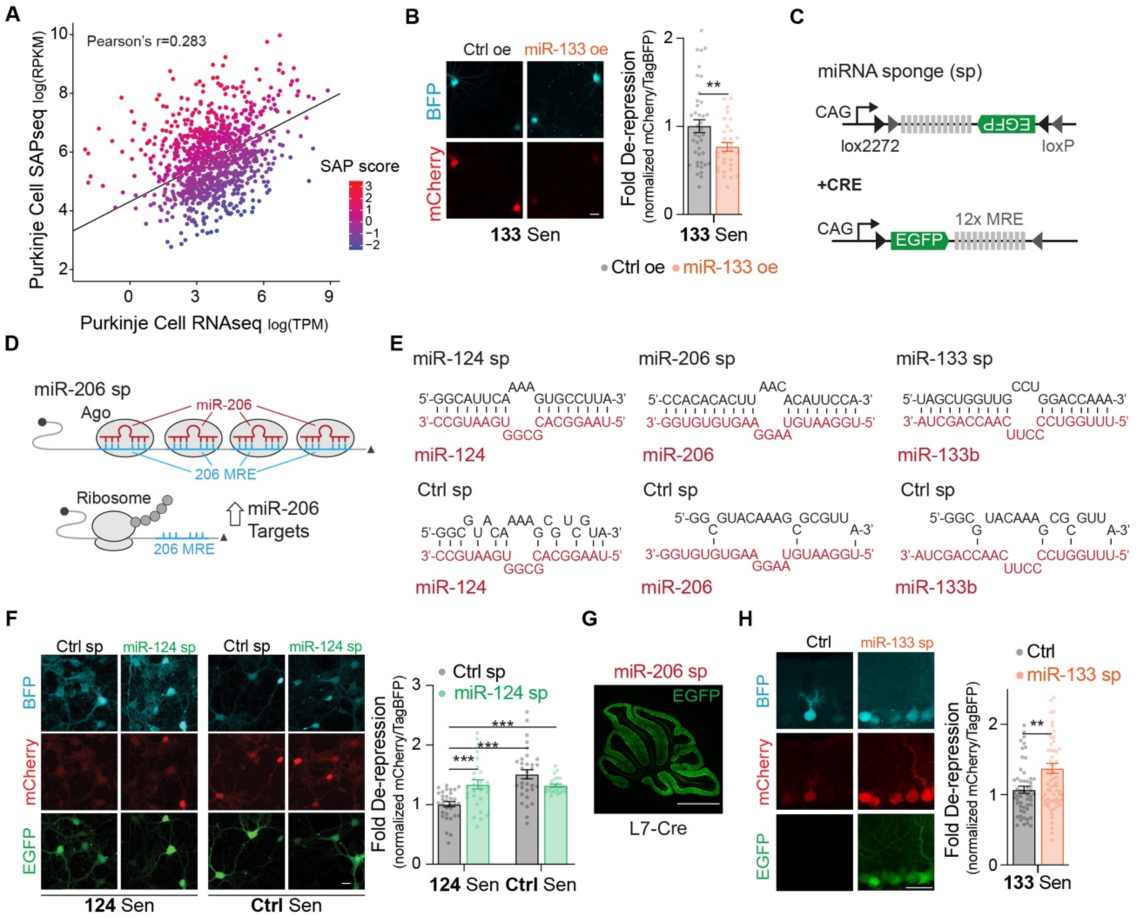

**Figure S6. Validation of the miRNA sponges, related to Figure 6**
(A) SAP score for P15 PCs miRNA targets. The line represents best-fit Deming regression. The SAP score for each target is calculated based on distance from the regression line.
(B) Validation of miR-133b overexpression (oe) in PNs with a miR-133b sensor (133 Sen). Scale bar, 10 µm. N=31-40 cells.
(C) Schematic of Cre-dependent miRNA sponge AAV constructs. The sponge consists of a GFP with an artificial 3' UTR containing 12 high-affinity miRNA binding sites.
(D) Schematic of miR-206 sponge activity (miR-206 sp). The sponge binds and sequesters miR-206, inducing its loss-of-function.
(E) Sequences and RNA duplexes of the miR-124, miR-206, miR-133 and Ctrl sponges (miR-124 sp, miR-206 sp, miR-133 sp, Ctrl sp).
(F) Validation of the miR-124 sponge in cultured cortical PNs with the miR-124 sensor (124 Sen) at DIV8. The sponge was induced with an AAV expressing Cre under the neuron-specific hSyn promoter. miR-124 sp de-repressed mCherry. Scale bar, 10 µm. N=24-32 cells.
(G) Sagittal section of an $L7^{Cre}$ mouse cerebellum transduced with the miR-206 sp at P28. The sponge is only expressed in PCs.
(H) Validation of the miR-133 sponge with 133 Sen in P28 PCs. miR-133 sp de-repressed mCherry. The Ctrl here are littermate wild-type mice ($L7^{Cre}$ is a BAC line) transduced with 133 Sen. Scale bar, 50 µm. N=54, 71 cells from 3 mice.
Data are mean ± SEM. Statistics for (A): Deming regression; (B), (F) and (H): Welch's t-test. $**p \leq 0.01$; $***p \leq 0.001$.
